# PIK3CA in *Kras^G12D^/Trp53^R172H^* Tumor Cells Promotes Immune Evasion by Limiting Infiltration of T Cells in a Model of Pancreatic Cancer

**DOI:** 10.1101/521831

**Authors:** Nithya Sivaram, Patrick A. McLaughlin, Han V. Han, Oleksi Petrenko, Ya-Ping Jiang, Lisa M. Ballou, Kien Pham, Chen Liu, Adrianus W.M. van der Velden, Richard Z. Lin

**Affiliations:** Department of Physiology and Biophysics, Stony Brook University, Stony Brook, N.Y.; Molecular and Cellular Biology Graduate Program, Stony Brook University, Stony Brook, N.Y.; Department of Molecular Genetics and Microbiology and Center for Infectious Diseases, Stony Brook University, Stony Brook, N.Y.; Department of Pathology and Laboratory Medicine, New Jersey Medical School and Robert Wood Johnson Medical School, Rutgers University School of Medicine, Newark, N.J.; Medical Service, Northport VA Medical Center, Northport, N.Y.

## Abstract

The presence of tumor-infiltrating T cells is associated with favorable patient outcomes, yet most pancreatic cancers are immunologically silent and resistant to currently available immunotherapies. Here we show using a syngeneic orthotopic implantation model of pancreatic cancer that *Pik3ca* regulates tumor immunogenicity. Genetic silencing of *Pik3ca* in *Kras^G12D^/Trp53^R172H^*-driven pancreatic tumors leads to infiltration of T cells, complete tumor regression, and 100% survival of immunocompetent host mice. By contrast, *Pik3ca*-null tumors implanted in T cell-deficient mice progress and kill all of the animals. Adoptive transfer of tumor antigen-experienced T cells eliminates *Pik3ca*-null tumors in immunodeficient mice. Loss of PIK3CA or inhibition of its effector, AKT, increases the expression of MHC Class I and CD80 on tumor cells. These changes contribute to the increased susceptibility of *Pik3ca*-null tumors to T cell surveillance. These results indicate that tumor cell PIK3CA-AKT signaling limits T cell recognition and clearance of pancreatic cancer cells. Strategies that target this pathway may yield an effective immunotherapy for this cancer.

**SIGNIFICANCE:** PIK3CA-AKT signaling in pancreatic cancer cells limits T cell infiltration and clearance of tumors by suppressing the surface expression of MHC Class I and CD80. Targeting the PIK3CA-AKT pathway in tumor cells provides a new avenue for discovery of novel pancreatic cancer immunotherapies.

## INTRODUCTION

Pancreatic ductal adenocarcinoma (PDAC) is the third leading cause of cancer-related death in the USA, with 44,330 deaths in 2018 and a five-year survival rate of only 8.5% [1]. Standard chemotherapies have little impact on PDAC patient survival, and even those patients who are suitable for surgical resection have only a 10% survival rate past five years [2]. Presence of tumor-infiltrating T cells is correlated with a better prognosis and increased survival of PDAC patients [3-6]. Recognition of tumor cells by cytotoxic CD8^+^ T cells requires binding of the T cell receptor (TCR) to antigens presented on major histocompatibility complex class I (MHC I), which consists of a membrane-spanning heavy chain and β2 microglobulin (B2m). MHC I is expressed on the surface of all nucleated cells. Pancreatic cancer cell lines isolated from *Kras^G12D^* or *Kras^G12D^;Trp53^R172H^* mice [7] and many human pancreatic cancers and pancreatic cancer cell lines [8, 9] have low MHC I levels that might contribute to immune evasion. In addition to TCR activation, the co-receptor CD28 expressed on T cells is stimulated by CD80. Without CD80, activated T cells become anergic even in the presence of a presented antigen. CD80 is expressed mainly on the surface of professional antigen-presenting cells (APCs) but is sometimes detected at low levels on tumor cells. Upregulation of CD80 on tumor cells has been shown to render them more susceptible to lysis by T cells [10-12].

More than 90% of PDACs have oncogenic mutations in the *KRAS* gene [13]. Phosphoinositide 3-kinase (PI3K) is a critical downstream effector of KRAS. Class I PI3Ks are heterodimeric lipid kinases consisting of a regulatory subunit bound to one of four different p110 catalytic subunits (PIK3CA (also called p110α), PIK3CB, PIK3CD or PIK3CG). In normal cells, PI3Ks regulate proliferation, survival, and differentiation. Upon activation, PI3Ks catalyze the conversion of phosphatidylinositol-4,5-bisphosphate to phosphatidylinositol-3,4,5-trisphosphate (PIP_3_). PIP_3_ then recruits downstream effectors such as the protein kinase AKT to the cell membrane, aiding in its phosphorylation and activation [14]. The PI3K-AKT signaling cascade is among the most frequently dysregulated and hence extensively studied pathways in human cancers [15-17].

PI3Ks in pancreatic cells and in immune cells have been shown to affect pancreatic tumorigenesis and cancer growth. Oncogenic KRAS^G12D^ signals through PIK3CA, but not PIK3CB, to induce acinar-to-ductal metaplasia that is required for pancreatic tumor formation [18], and the catalytic activity of PIK3CA is required for this tumorigenic process [19]. It is not yet known if PIK3CA is also required for the growth and progression of established PDAC *in vivo.* Moreover, PIK3CA’s role in cancer immune surveillance has not been studied. Signaling by PIK3CG and PIK3CD in leukocytes, but not in tumor cells, can affect how the immune system responds to tumors in murine models, including pancreatic cancer [20, 21]. Small molecules that inhibit all PI3K isoforms have little effect or transient suppressive effects on PDAC growth in mouse models (reviewed in [22] and [23]). However, these drugs inhibit PI3Ks in both the tumor cells and immune cells, so the results of systemic inhibitor studies should be interpreted with caution.

In this study, we examined the function of PIK3CA in a pancreatic cancer cell line derived from mice expressing KRAS^G12D^ and TRP53^R172H^ [24, 25]. We show that cells lacking PIK3CA are viable *in vitro* but are cleared by tumor-infiltrating T cells when implanted in the pancreas of immunocompetent mice. The susceptibility of these tumors to immunosurveillance is due at least in part to increased expression of MHC I and CD80 on the tumor cell surface.

## RESULTS

### *Pik3ca* in Implanted KPC Cells Promotes Tumor Progression and Lethality

Both *Pik3ca* and *Egfr* have been shown to be required for KRAS^G12D^-induced pancreatic tumorigenesis [18, 19, 26]. To investigate the possible roles of *Pik3ca* and *Egfr* in pancreatic cancer progression, we used the FC1245 pancreatic cancer cell line that was isolated from a *Kras^LSL-G12D/+^;Trp53^LSL-R172H/+^;Pdx1-Cre* mouse in the C57BL/6 (B6) genetic background [25].

We first produced a parental cell line that stably expresses luciferase (referred to as wildtype (WT) KPC). We extracted genomic DNA from WT KPC cells to sequence exon 1 of the *Kras* gene and confirmed that the cells have a G-D mutation at codon 12. We did not detect the WT allele of the *Kras* gene (Supplementary Fig. S1). Such loss of the WT *Kras* allele is common in cell lines derived from both genetically engineered mouse models and patients with PDAC and other cancers [27-30]. Loss of WT KRAS is frequently associated with amplification in the mutant allele and has been shown to increase the aggressiveness and migration of PDAC and other cancers [27-31]. We then used CRISPR/Cas9 to produce KPC cell lines that lack either *Pik3ca* or *Egfr* (referred to as αKO or EgfrKO, respectively). Complete loss of PIK3CA or EGFR was confirmed by western blotting (Supplementary Fig. S2A). Immunoblotting and reverse phase protein array (RPPA) analysis also revealed changes in signaling due to ablation of *Pik3ca* or *Egfr*. In particular, there was a large decrease in AKT phosphorylation at S473 and T308 in αKO cells as compared to WT cells, while EgfrKO cells had higher levels of AKT phosphorylation than WT cells (Supplementary Fig. S2). WT and EgfrKO KPC cells proliferated at similar rates in standard 2D culture conditions, whereas αKO cells proliferated at about half the rate of WT cells (Fig. 1A). The percentage of cells stained with annexin V in each culture was not significantly different, indicating that the decreased proliferation rate of αKO KPC cells is not due to increased apoptosis (Fig. 1B). When grown under 3D culture conditions, all three cell lines formed compact spheroid colonies (Fig. 1C), indicating a capacity for anchorage-independent growth.

**Fig. 1.**
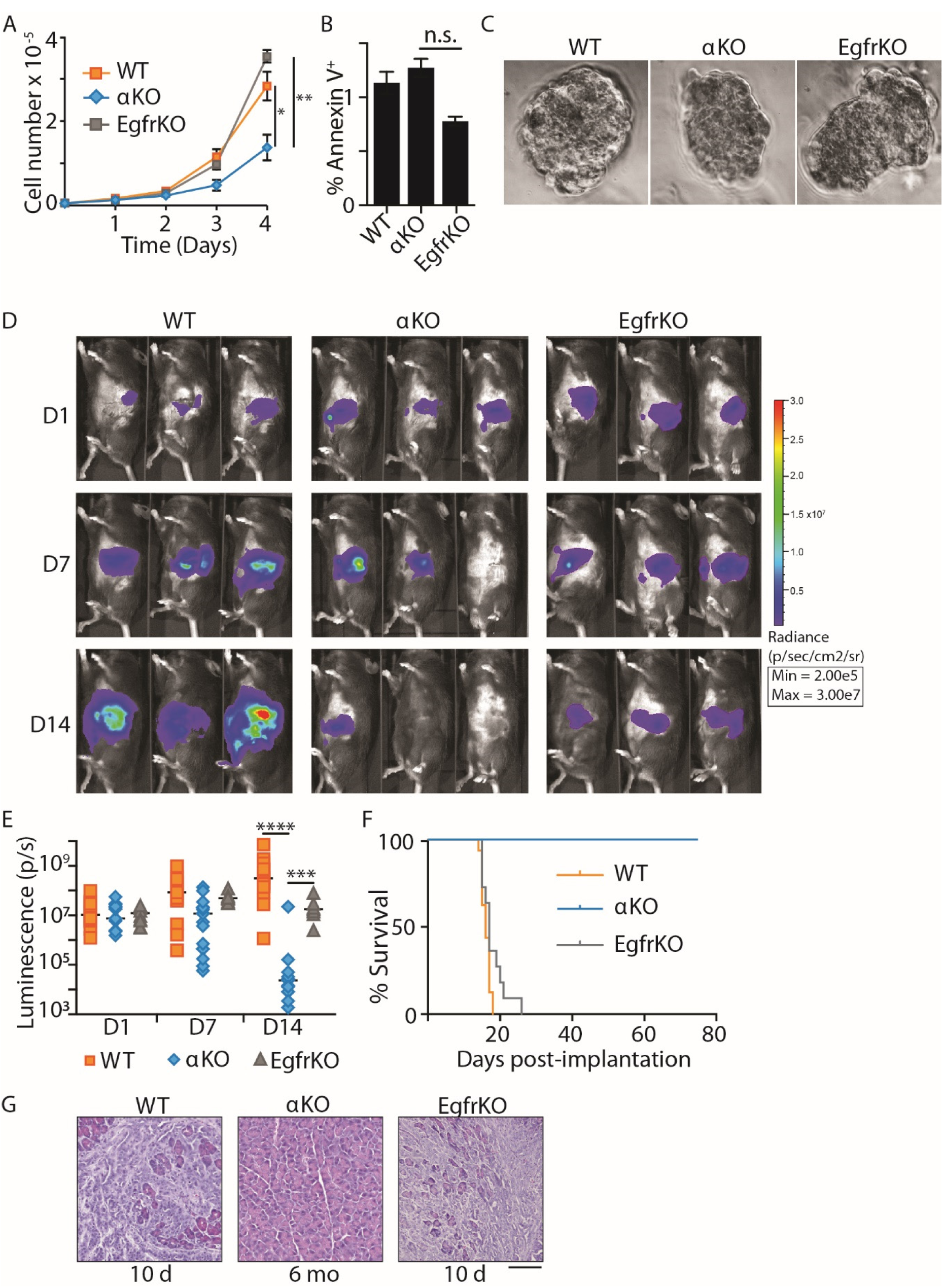
Proliferation of αKO pancreatic cancer cells *in vitro* and regression *in vivo*. **(A)** Proliferation rates of KPC cell lines in standard 2D culture. Cells plated in triplicate were counted at the times indicated (mean ± SEM; *n* = 3). \*\**P* = 0.033, WT *vs*. αKO, n.s., not significant (Student’s *t*-test). **(B)** Percentage of cells positive for annexin V staining at the 4-day time point in **A** (mean ± SEM, *n* = 3; n.s., not significant). **(C)** Representative light microscopy images (40X magnification) of spheroids formed after 5 days in 3D methylcellulose culture. **(D)** Cells (0.5 million) were implanted in the head of the pancreas of B6 mice. Tumor growth was monitored by IVIS imaging of the luciferase signal on days 1, 7, and 14 after implantation. Representative images of 3 mice in each group are shown. **(E)** Quantification of luciferase signals from each mouse. The bars indicate median. \*\*\*\**P* < 0.0001, WT vs. αKO, \*\*\**P* < 0.0005, WT vs. EgfrKO (Mann-Whitney U test). **(F)** Kaplan-Meier survival curves for B6 mice implanted with the indicated cell lines. Median survival: WT, 16 days (*n* = 16); EgfrKO, 17 days (*n* = 10); all αKO mice were alive at day 80 (*n* = 17). *P* < 0.0001, WT vs. αKO (log-rank test). **(G)** H&E-stained sections of pancreatic tissue collected at the indicated times after cell implantation in B6 mice. Scale bar,100 μm.

We next studied the growth of these cell lines *in vivo*. WT, αΚΟ, or EgfrKO cells were implanted in the head of the pancreas of syngeneic B6 mice, and tumor growth was monitored longitudinally by IVIS imaging. As expected, WT cells grew rapidly and formed large tumors (Fig. 1D). Median tumor volume quantified as total luminescence flux showed a 29.6-fold increase from day 1 to day 14 (Fig. 1E). Tumors formed by EgfrKO cells grew at a significantly slower rate (Fig. 1D and E). Mice implanted with WT or EgfrKO cells died 2 to 3 weeks after implantation, with a median survival of 16 days or 17 days, respectively (Fig. 1F). In contrast, tumors formed by implanted αKO cells showed an increase in median size from day 1 to day 7, and then the tumors regressed so that the luciferase signal was undetectable in 16 of 17 mice on day 14 (Fig. 1D and E). By day 21, the luciferase signal was undetectable in all 17 mice (data not shown). All B6 mice implanted with αKO cells were alive 80 days later (Fig. 1F). Some of these convalescent animals have been kept for up to 18 months after tumor implantation without any overt signs of illness. Implantation of B6 mice with another clone of αKO cells yielded similar results (Supplementary Fig. S3). At necropsy, mice implanted with WT or EgfrKO cells exhibited large pancreatic tumors and metastases to the peritoneum, liver, diaphragm, and lungs. H&E staining of pancreatic sections confirmed the presence of large tumors with a strong desmoplastic response (Fig. 1G). Some of the mice implanted with αKO cells were euthanized after 6 months: no abnormal lesions in any of the organs were seen by visual inspection, and no tumors were found in pancreatic sections of these animals (Fig. 1G).

### Regression of αKO KPC Tumors is Linked to T Cell Infiltration

To investigate if a host immune response might be responsible for the regression of αKO tumors in our mouse model, we implanted WT, αKO, or EgfrKO tumor cells in the pancreas of B6 mice and sacrificed the animals 10 days later. H&E staining of pancreatic sections from all three groups showed substantial areas of normal pancreas interspersed with tumors (Fig. 2A). Immunohistochemistry showed that αKO tumors were infiltrated with large numbers of CD3^+^, CD4^+^, and CD8^+^ cells (e.g., T cells) (Fig. 2A and B). Similar to most human pancreatic tumors, WT and EgfrKO tumors were associated with peri-tumoral T cells, but few T cells were seen infiltrating the tumors (Fig. 2A and B). We did not observe a significant difference in the number of F4/80^+^ macrophages infiltrating the tumors in the three groups (Fig. 2A and B).

**Fig. 2.**
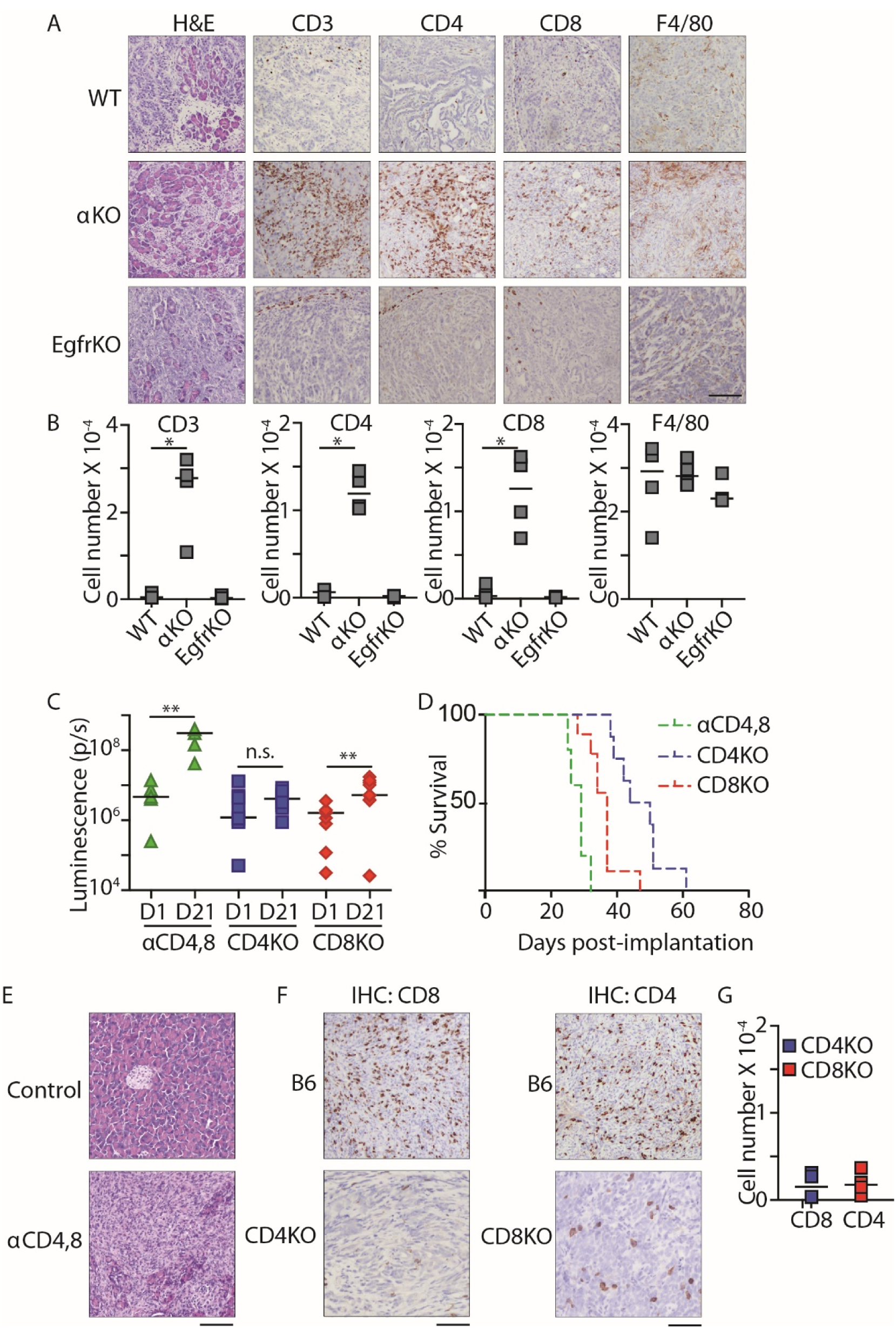
T cell infiltration of αKO tumors and the requirement of CD4^+^ and CD8^+^ T cells for αKO tumor regression. **(A and B)** WT, αKO or EgfrKO cells (0.5 million) were implanted in the head of the pancreas of B6 mice, and pancreata were harvested 10 days later (EgfrKO, *n* = 3; WT and αKO, *n* = 4). **(A)** Sections were stained with H&E, or IHC was performed with the indicated antibodies. Representative sections are shown. Scale bar, 100 μm. **(B)** Quantification of tumor-infiltrating cells positive for CD3, CD4, CD8, or F4/80. \**P* = 0.0286 (Mann-Whitney U test). **(C-F) α**KO cells (0.5 million) were implanted in the head of the pancreas of B6 mice injected with neutralizing CD4 and CD8 antibodies (αCD4,8, green), CD4KO (blue), or CD8KO (red) mice. **(C)** Quantification of luciferase signals from IVIS images of each mouse. Bars indicate median. \*\**P*< 0.005 and n.s., not significant (two-tailed Wilcoxon signed-rank test). **(D)** Kaplan-Meier survival curves for αCD4,8 (*n* = 5, median survial: 29 days), CD4KO (*n* = 9, median survival: 47 days) and CD8KO (*n* = 8, median survival: 37 days) mice implanted with αKO cells. *P* = 0.0001 (log-rank test). **(E)** H&E staining of pancreatic sections collected at death of αCD4,8 mice vs. saline injected control mouse. **(F)** IHC staining of pancreatic sections with antibodies against CD4 or CD8. Pancreata were collected from CD4KO or CD8KO mice at the humane endpoint. For comparison, sections of pancreata collected 10 days after implantation of B6 mice with 0.5 million αKO cells are shown. Scale bars, 100 μm. **(G)** Quantification of tumor-infiltrating CD4^+^ or CD8^+^ T cells from pancreatic sections in F.

To test if tumor-infiltrating T cells are responsible for tumor regression, we injected B6 mice with neutralizing antibodies to deplete CD4^+^ and CD8^+^ T cells. Depletion of T cells was confirmed by flow cytometry (Supplementary Fig. S4A). We then implanted these mice with αKO cells. IVIS imaging showed that αKO tumors grew in these mice. The average fold-change in tumor volume from day 1 to day 21 was 65.7 (Fig. 2C). All mice that received neutralizing antibodies died due to tumor growth (Fig. 2D and E). As expected, the control mouse that did not receive injections of neutralizing antibodies showed complete tumor regression (Fig. 2E). To determine the specific subtype of T cells involved in tumor regression, we implanted αKO cells in immunocompromised mice lacking CD4^+^ or CD8^+^ T cells (referred to as CD4KO and CD8KO). αKO tumors did not regress in CD4KO or CD8KO mice and all of the animals died from tumor growth (Fig. 2C and D and Supplementary Fig. S4B). Tumor growth was slower in the CD4KO mice, and they survived longer than the CD8KO animals (median survival of 47 days *vs*. 37 days, respectively). Further, CD4KO and CD8KO mice survived significantly longer than B6 mice depleted of CD4^+^ and CD8^+^ T cells by neutralizing antibodies (median survival of 29 days) (Fig. 2D). Immunohistochemistry showed a decrease in the number of tumor-infiltrating CD8^+^ T cells in CD4KO mice and fewer tumor-infiltrating CD4^+^ T cells in CD8KO mice as compared to tumors in B6 animals (Fig. 2E and F). These results suggest that both CD4^+^ and CD8^+^ T cells are essential and that they cooperate with each other to infiltrate αKO tumors and cause tumor regression.

### Adoptive T Cell Transfer Protects SCID Mice from αKO KPC Tumors

We next investigated if T cells from a convalescent B6 mouse (implanted with αKO cells and recovered for 6 months; see Fig. 1G) can target and kill αKO cells implanted in mice with severe combined immune deficiency (SCID) that lack functional T and B cells. We first implanted αKO cells in the head of the pancreas of a control group of SCID mice. The tumors grew rapidly and killed all of the animals (Fig. 3A-C). A second group of SCID mice was implanted with αKO cells one day after receiving an adoptive transfer of CD90.2^+^ T cells enriched from the spleens of convalescent mice. The αKO tumors regressed in these animals (Fig. 3A and B) and all of the mice survived for >80 days without overt signs of illness (Fig. 3C). H&E-stained pancreatic sections showed that the control group of SCID mice had large tumors with little normal tissue, while mice that received T cells had completely normal pancreas 4 months after implantation (Fig. 3D).

**Fig. 3.**
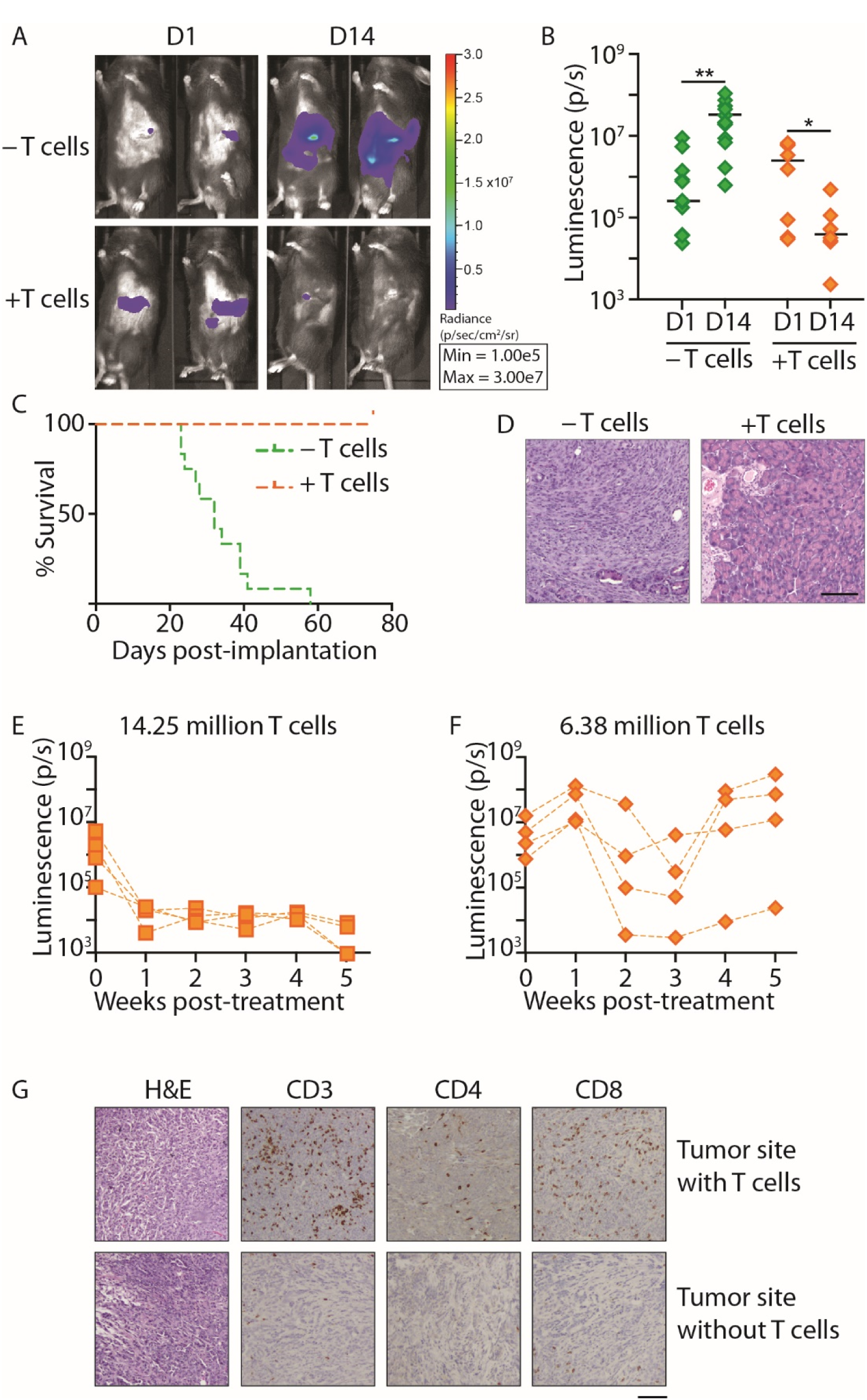
Adoptive T cell transfer protects SCID mice from implanted αKO tumors. **(A-D)** SCID mice received either no pretreatment (-T cells, *n* = 12) (green) or adoptive transfer of 5 million T cells harvested from B6 mice previously implanted with αKO cells (+T cells, *n* = 8) (orange). One day later, αKO cells (0.5 million) were implanted in the head of the pancreas of each animal. **(A)** Tumor growth was monitored by IVIS imaging. Representative images of 2 mice in each group are shown. **(B)** Quantification of luciferase signals from each mouse. Bars indicate median. \**P* = 0.0156 and \*\**P* = 0.0024 (Wilcoxon signed-rank test). **(C)** Kaplan-Meier survival curves. Median survival: –T cells, 32 days; +T cells, all were alive at day 80. *P* < 0.00001 (log-rank test). **(D)** Representative H&E-stained pancreatic sections. Pancreata were collected at the humane endpoint (-T cells) or 4 months after tumor implantation (+T cells). Scale bar, 100 μm. **(E-G)** 0.5 million αKO KPC cells were implanted in the head of the pancreas of SCID mice (*n* = 8). Four days later (t = 0), the mice were imaged by IVIS and then treated with 14.25 million or 6.38 million T cells from a convalescent B6 mouse previously implanted with αKO cells (*n* = 4 per group). Tumor growth was monitored by IVIS imaging. **(E** and **F)** Quantification of luciferase signals over time. **(G)** Mice treated with 6.38 million T cells were sacrificed at week 5 and their pancreata were collected for histology. Representative H&E and IHC images of different tumor sites from the same mouse are shown. Scale bar, 100 μm.

We next asked if adoptive T cell transfer is an effective treatment for established αKO tumors. Two groups of SCID mice were implanted with αKO cells in the pancreas. After determining tumor size 4 days later by IVIS imaging (Fig. 3E and F, t = 0), one group of animals was given 14.25 million T cells from convalescent mice and the second group was treated with 6.38 million T cells. Mice that received 14.25 million T cells showed complete tumor regression after one week (Fig. 3E) and all of the animals survived for >10 weeks. In mice that received fewer T cells, the tumors regressed but started growing again after 3-4 weeks (Fig. 3F). We collected pancreata from the animals treated for 5 weeks with 6.38 million T cells and examined pancreatic sections by immunohistochemistry. Some of the αKO tumors were infiltrated by CD4^+^ and CD8^+^ T cells (*top panels*, Fig. 3G), whereas others from the same mouse were scarcely infiltrated with T cells (*bottom panels*, Fig. 3G). These results suggest that T cells from mice previously exposed to the αKO tumor can be used to induce tumor regression in another animal as long as the number of T cells relative to tumor cells is adequate. Indeed, the significance of T cell number and the time of adoptive T cell transfer to eradicate tumors has been reported [32].

### *Pik3ca*-dependent Suppression of MHC I and CD80 Levels Contributes to Immune Evasion of WT KPC Tumors

We first analyzed RNAseqV2 and RPPA data from The Cancer Genome Atlas (TCGA) to check for a correlation between the effectors of the PI3K signaling pathway and CD80 and MHC I. We assessed the mRNA levels of CD80 and B2m in 180 PDAC patients and observed a positive correlation with PTEN mRNA expression. Patients that showed high PTEN expression also had high levels of B2m and CD80 mRNA. Conversely, patients with low PTEN mRNA had low mRNA levels of CD80 and B2m (Fig. 5A and B). In addition, analysis of the RPPA data showed that pAKT S473 expression was inversely correlated with B2m mRNA levels in PDAC patients (Fig. 5C and D). Our analyses suggest that PI3K signaling in the tumor cells regulates the expression of MHC I and CD80 in PDAC.

We wondered if a difference in expression of MHC I and CD80 might explain the different host immune responses to implanted αKO or WT KPC cells. Indeed, flow cytometry showed that cell surface expression of the MHC I heavy chain (H-2K^b^ in B6 mice) was 6.4 times higher in αKO KPC cells than in WT cells (Fig. 4A; average geometric means of 14.7 *vs.* 2.3, respectively). We also found that cell surface CD80 was 4.4 times higher in αKO cells (Fig. 4B; average geometric mean, 36.8 *vs*. 8.4, respectively). H-2K^b^ and B2m mRNA levels tended to be higher in αKO cells, but the differences were not statistically significant. By contrast, CD80 mRNA levels were significantly higher in αKO cells (Fig. 4C). Priming of CD4^+^ helper T cells requires binding of the TCR to antigens presented on MHC Class II (MHC II), which is normally expressed in immune cells, but it has also been detected on some cancer cells [33, 34]. However, cell surface expression of MHC II was not detected in either cell line (Supplementary Fig. S5A). These results suggest that *Pik3ca*-dependent suppression of MHC I and CD80 in WT KPC cells may allow the tumors to evade lysis by CD8^+^ T cells. CD4^+^ T cells may provide helper functions to the CD8^+^ T cells that infiltrate αKO tumors.

**Fig. 4.**
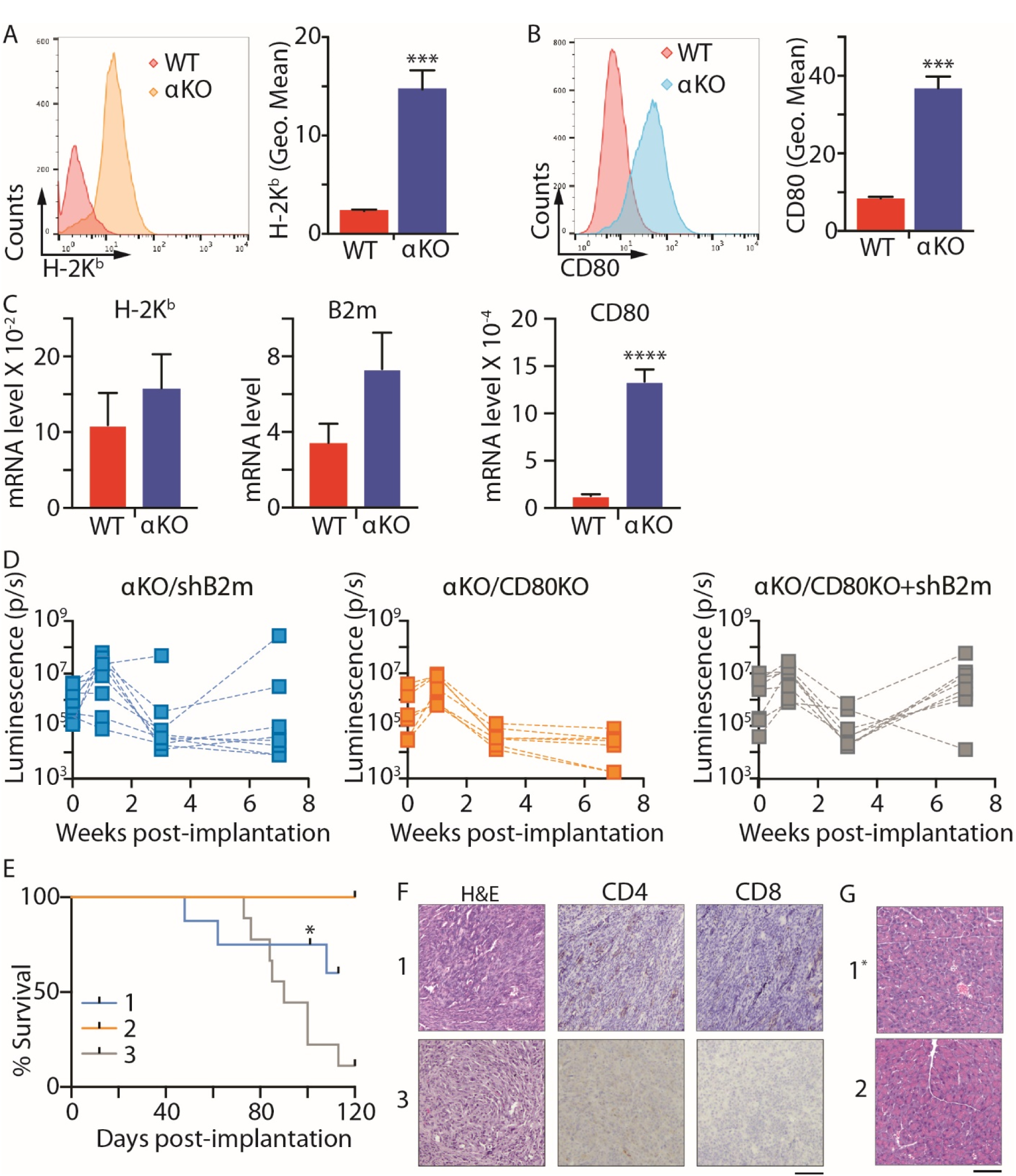
PIK3CA regulates cell surface expression of MHC I and CD80 in KPC cells. Flow cytometric analysis of cell surface levels of H-2K^b^ **(A)** and CD80 **(B)** in WT and αKO cells. The *left panels* show representative histograms and the *right graphs* show the mean ± SEM of the geometric means (Geo. Mean) of each flow cytometry distribution. H-2K^b^, *n* = 4; CD80, *n* = 3. **\*\***\**P* = 0.0007 (unpaired *t-*test) **(C)** qRT-PCR analysis of mRNA expression of H-2K^b^, B2m, and CD80 in WT and αKO cells. Gene expression changes were normalized to *Hprt*. Graph shows mean±SEM (*n* = 6). \*\*\*\**P* < 0.0001 (unpaired *t*-test) **(D-G)** B6 mice were implanted with 0.5 million αKO/shB2m, αKO/CD80KO, or αKO/CD80KO+shB2m cells in the head of the pancreas and tumor growth was monitored by IVIS imaging. **(D)** Quantification of the luciferase signals in each mouse. αKO/shB2m, *n* = 8; αKO/CD80KO, *n* = 7; αKO/CD80KO+shB2m, *n* = 8. **(E)** Kaplan-Meier survival curves. 1, αKO/shB2m; 2, αKO/CD80KO; 3, αKO/CD80KO+shB2m; *, one mouse was euthanized for histology and removed from the study. Median survival: αKO/shB2m, 110.5 days; αKO/CD80KO+shB2m, 87.5 days. All αKO/CD80KO mice were alive at 115 days. *P* = 0.0026 (log-rank test). **(F)** Representative H&E and IHC images of pancreatic sections from (1) αKO/shB2m or (3) αKO/CD80KO+shB2m mice that died of tumor progression. Scale bar, 100 μm. **(G)** Representative H&E-stained pancreatic sections from (1) one αKO/shB2m mouse or (2) α/CD80KO mice euthanized at 101 or 160 days, respectively. Scale bar, 100 μm.

The roles of MHC I and CD80 in immune-mediated regression of αKO tumors were furthered investigated using αKO cell lines with reduced expression of these proteins. CRISPR/Cas9 was used to target *Cd80* in αKO cells to generate αKO/CD80KO cells. MHC I cannot be knocked out completely in cells used for implantation experiments because natural killer cells recognize and kill cells that lack the self MHC I. Therefore, we used shRNA to knock down the level of B2m in αKO or αKO/CD80KO cells to produce αKO/shB2m or αKO/CD80KO+shB2m cell lines, respectively. The new αKO cells expressed B2m and CD80 at levels comparable to the WT cells. Flow cytometry confirmed a reduction in CD80 and/or H-2K^b^ on the cell surface in the appropriate cell lines (Supplementary Fig. S5B). We observed no significant difference in proliferation rate of the three cell lines in 2D culture as compared to αKO cells (Supplementary Fig. S5C). Implantation of αKO/shB2m cells in the pancreas of B6 mice resulted in initial tumor growth in all of the animals over the first week. By week 7, however, the tumors had regressed in 5 of 8 mice (Fig. 4D and Supplementary Fig. S6), and these 5 mice lived for more than 100 days (Fig. 4E). Three mice died of tumor progression, and necropsy showed a solid mass in the pancreas and metastases to the peritoneum, diaphragm, liver, and lungs. Implanted αKO/CD80KO tumors grew larger over the first week and then regressed in every animal (Fig. 4D and Supplementary Fig. S6). All of the animals implanted with αKO/CD80KO cells survived >100 days (Fig. 4E). In contrast, 7 of 8 mice implanted with αKO/CD80KO+shB2m cells still had detectable pancreatic tumors at week 7 (Fig. 4D and Supplementary Fig. S6), and all these animals eventually died from tumor progression (Fig. 4E). Examination of pancreatic tissue from αKO/shB2m or αKO/CD80KO+shB2m mice that succumbed to tumor progression showed large tumors that were devoid of infiltrating CD4^+^ or CD8^+^ T cells (Fig. 4F). In summary, αKO cells with high surface CD80 and MHC I regressed when implanted in B6 mice (Figure 1D-F). In contrast, αKO cells with low surface B2m alone (αKO/shB2m) or CD80 alone (αKO/CD80KO) were eliminated in most of the animals (Figure 4D). However, αKO cells with decreased surface expression of both B2m and CD80 (αKO/CD80KO+shB2m) were not cleared by the T cells and resulted in the death of the mice (Figure 4E) similar to WT cells (Figure 1D-F). No tumors were detected in pancreatic tissue from one αKO/shB2m mouse or αKO/CD80KO mice that were sacrificed at 101 days or 160 days, respectively (Fig. 4G). These results indicate that upregulation of both MHC I and CD80 contribute to immune evasion by αKO KPC tumors. The extended survival of mice with αKO/CD80KO+shB2m tumors as opposed to WT KPC tumors suggests that other tumor-specific or host-related factors also play a role in tumor progression.

### AKT Acting Downstream of PIK3CA is Responsible for Immune Evasion of KPC cells

Because AKT activity is significantly higher in WT KPC cells than in αKO cells (Supplementary Fig. S2), we wondered if AKT acts downstream of PIK3CA to suppress MHC I and CD80 levels. Flow cytometry of WT cells treated with increasing concentrations of an AKT inhibitor (Akti) showed a dose-dependent increase in expression of H-2K^b^ and CD80 (Fig. 5A). The increases in average geometric mean at 20 μM Akti *vs.* DMSO were 5.5-fold (H-2K^b^) and 4.4-fold (CD80). These increases in expression are comparable to the different expression levels in αKO *vs.* WT KPC cells (6.3-fold (H-2K^b^) and 4.4-fold (CD80); Fig. 4A and B). Akti treatment also significantly increased the cell surface expression of both MHC I (HLA-ABC in human) and CD80 in 3 of 8 human PDAC cell lines (Fig. 5B). This result is not surprising since there are potentially many mechanisms by which MHC I can be downregulated. For example, cancer cells with biallelic loss of MHC I will not respond to Akt inhibition. Somewhat surprisingly, all of the human PDAC cell lines tested showed upregulated CD80 expression after Akti treatment (Fig. 5B). Western blotting of the mouse and human PDAC cell lines showed that Akti treatment reduced the level of phospho-AKT (Supplementary Fig. S7).

**Fig. 5.**
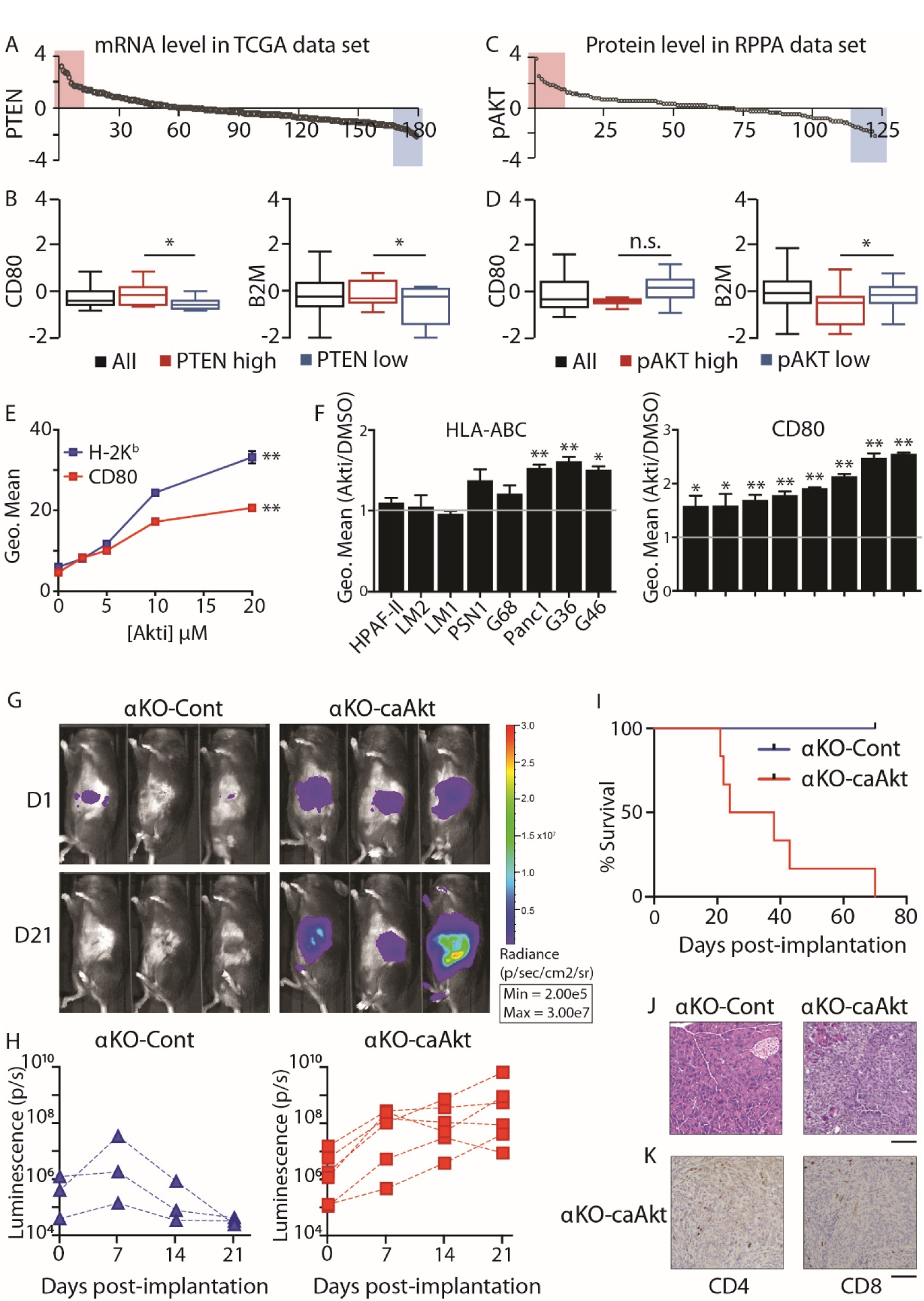
PIK3CA regulation of CD80 and MHC I is mediated by AKT signaling. **(A)** PTEN mRNA distribution in pancreatic cancer sections from the TCGA database. **(B)** Box plots showing CD80 and B2M mRNA expression in patients with high and low PTEN expression. **(C)** Phospho-Akt (pAKT) levels in pancreatic cancer sections from the TCGA database. **(D)** Box plots showing CD80 and B2M mRNA expression in patients with high and low pAKT levels. *N* = 10, \**P* < 0.05, *n.s.* not significant. **(E)** WT KPC cells were treated with increasing concentrations of Akt inhibitor (Akti) for 48 hours. Cell surface levels of CD80 and H-2K^b^ were quantified by flow cytometry. The mean ± SEM of the geometric means of each flow cytometry distribution is plotted *vs*. inhibitor concentration. *n* = 3 for each inhibitor concentration. \*\**P* = 0.0031 (H-2K^b^) and \*\**P* = 0.0018 (CD80) for DMSO *vs.* 20 μM Akti (paired *t*-test). **(F)** Human pancreatic cell lines were treated with 10 μM Akti for 48 hours. Cell surface levels of HLA-ABC *(left)* and CD80 *(right)* were determined by flow cytometry. Each treatment group was normalized to DMSO (*n* = 3). The graph shows fold change (mean ± SEM) over DMSO. \**P* < 0.05, \*\**P* < 0.005. **(G-K) αΚΟ**-Cont or αKO-caAkt cells (0.5 million) were implanted in the head of the pancreas of B6 mice. **(G)** Representative IVIS images showing tumor size on days 1 and 21. **(H)** Quantification of luciferase signals in each mouse. αΚΟ-Cont, *n* = 3 and αKO-caAkt, *n* = 6.**(I)** Kaplan-Meier survival curves. Median survival: αKO-caAkt, 31 days. All αKO-Cont mice were alive at 71 days. *P* = 0.011 (log-rank test). **(J)** Representative H&E-stained pancreatic sections from mice implanted with αΚΟ-Cont cells (euthanized at 71 days) or αKO-caAkt cells (post-mortem). **(K)** Representative IHC images of pancreatic sections from mice implanted with αKO-caAkt cells (post-mortem). For **J** and **K,** *n* = 3. Scale bar, 100 μm.

We next tested if increasing AKT activity in αKO pancreatic tumors allows them to evade immune surveillance. Western blotting showed that αKO cell lines stably expressing a constitutively active form of AKT (αKO-caAkt cells) have high levels of phospho-Akt as compared to vector control cells stably expressing the control vector (αKO-Cont cells; Supplementary Fig. S8A). αKO-caAkt cells proliferated about 1.4 times faster than αKO-Cont cells in 2D culture (Supplementary Fig. S8B). When implanted in the pancreas of B6 mice, the αKO-Cont tumors grew over the first 7 days and then regressed, and all of the mice lived for >70 days (Fig. 5C-E). In contrast, the αKO-caAkt cells grew rapidly, formed large tumors, and killed all of the animals (Fig. 5C-E). Mice that died from implanted αKO-caAkt cells had large pancreatic tumors and extensive metastases to the liver, lungs, and colon. Some mice also had metastases to the diaphragm and peritoneum. Pancreatic sections showed tumors devoid of intra-tumoral CD4^+^ or CD8^+^ T cells, although T cells were present in tissue surrounding the tumors (Fig. 5F and G). These results are similar to what we observed with WT KPC cells implanted in B6 mice (see Fig. 1D-F). Mice implanted with αKO-Cont cells were sacrificed 71 days after implantation. H&E-stained pancreatic sections showed no abnormalities (Fig. 5F). These results suggest that PIK3CA-AKT signaling mediates immune evasion in WT KPC cells.

## DISCUSSION

Pancreatic tumors in patients are well known for their ability to exclude T cells [35-37], and limited T cell infiltration is also a prominent feature of pancreatic tumors in *Kras^LSL-G12D/+^;Trp53^LSL-R172H/+^;Pdx1-Cre* mice [35, 38, 39]. The presence of tumor-infiltrating cytotoxic T cells is correlated with increased survival of pancreatic cancer patients [3-6]. The salient finding of this study is that PIK3CA-AKT signaling regulates the susceptibility of KPC pancreatic tumors to T cell anti-tumor responses that result in infiltration and elimination of the tumor. *Pik3ca*-null tumors contained significantly more tumor-infiltrating T cells than WT tumors, and the αKO tumors regressed only in mice with functional T cells.

MHC I is part of the machinery that presents processed antigenic peptides to cytotoxic CD8^+^ T cells. Suppression of cell surface MHC I is a well-documented mechanism used by some tumors to evade the immune system [8]. Conversely, high MHC I expression in triple negative breast tumors is correlated with increased tumor-infiltrating lymphocytes and better prognosis in patients [40]. One important phenotype caused by *Pik3ca* loss in KPC cells is upregulation of MHC I on the cell surface. Treatment of WT KPC cells or some human PDAC cell lines with an AKT inhibitor caused a similar change. To our knowledge, this is the first report that PIK3CA-AKT signaling negatively regulates MHC I in cancer cells. Earlier studies identified some other pathways that regulate MHC I. A human kinome shRNA interference-based screen was used to discover several kinases that positively or negatively regulate MHC I expression [41]. MAP2K1 (also called MEK1) and EGFR were validated as negative regulators by showing that treatment of several cancer cell lines with inhibitors of either enzyme increased mRNA levels and cell surface expression of HLA-A and B2m [41]. Cytokines such as IFN-γ secreted by T helper cells have been shown to upregulate MHC I expression *via* JAK/STAT signaling [42]. Interestingly, a key factor mediating HIV immune escape is the HIV-1 protein Nef, which acts through PI3K to downregulate trafficking of MHC I to the cell surface [43]. Further studies are needed to identify the mechanisms downstream of PIK3CA and AKT that are responsible for downregulating MHC I in KPC cells.

A second important phenotype caused by *Pik3ca* loss or AKT inhibition in pancreatic cancer cells is upregulation of CD80 on the cell surface. CD80 binds to CD28 on T cells and provides a costimulatory signal that is indispensable for T cell activation. Most tumors do not express CD80, but engineered expression of endogenous or exogenous CD80 on tumor cells has been shown to enhance anti-tumor T cell responses and promote tumor regression [12, 44, 45]. Cytotoxic T lymphocyte antigen-4 (CTLA-4) is another CD80 receptor expressed on T cells. CD80 ligation with CTLA-4, unlike CD28, initiates a suppressive signaling cascade that leads to T cell exhaustion [46]. Upregulating CD80 on tumor cells potentially can result in T cell exhaustion and augment tumor progression. However, our results show that αKO KPC tumors were efficiently cleared by the immune system, suggesting that inhibitory signals through CTLA-4 are minor. We observed elevated levels of PD-L1 and PD-L2 in αKO cells and in AKTi-treated WT cells (Supplementary Fig. S9). PD-L1 and PD-L2 expressed on tumor cells potentially can suppress T cell anti-tumor responses [47] but do not exert a dominant inhibitory effect in our model. Interestingly, a similar increase in PD-L1 and PD-L2 was observed in gastric cancer patients and correlated with increased T cell infiltration in the tumors and better prognosis [48]. Deleting CD80 was not sufficient to convert “hot” αKO KPC tumors with infiltrating T cells into “cold” tumors that exclude T cells. Knock-down of MHC I was more successful in this regard, but αKO/shB2m tumors progressed in only 3 of 8 implanted animals. The combined actions of MHC I downregulation plus CD80 silencing were required to reverse the highly immunogenic phenotype of αKO KPC tumors. Because the median survival of mice with αKO/CD80KO+shB2m tumors (87.5 days) was much longer than that of mice with WT KPC tumors (16 days), we suspect that additional *Pik3ca*-regulated immune modulators, perhaps including cytokines, contribute to tumor immunogenicity.

Our data using CD4KO and CD8KO mice indicate that both CD4^+^ and CD8^+^ T cells are essential for αKO KPC tumor regression. Without one subtype of T cell, the other subtype was less able to infiltrate the tumors. CD4^+^ T helper cells (Th) secrete cytokines and chemokines such as IFNγ and IL-2 that provide helper functions to B cells, APCs and the cytotoxic CD8^+^ T cells that ultimately lyse tumors [32, 49]. Especially in MHC II-negative tumors, CD4^+^ T cells are critical during the effector phase of the anti-tumor response. Mice depleted of CD4^+^ T cells prior to tumor challenge were unable to clear the tumor [50]. In the context of PDAC, IFNγ-secreting Th1 cells were shown to slow tumor growth by stimulating the proliferation of CD8^+^ cytotoxic T cells, while IL-5-secreting Th2 cells promoted tumor growth by inhibiting CD8^+^ cells [51]. CD4^+^ T cells may also acquire cytotoxic activity *in vivo* as seen in some melanomas [52, 53], and can play a role in generating and maintaining memory CD8^+^ T cells [54, 55]. Adoptive T cell transfers of both CD4^+^ and CD8^+^ T cells have been shown to be more effective than either subtype alone [32, 56]. CD4^+^ and CD8^+^ T cell infiltrate in PDAC patient tumors is primarily of the effector memory type with few naïve T cells [37]. Because CD4^+^ T cells can alter the tumor microenvironment and facilitate CD8^+^ T cell trafficking and function at the tumor site [32, 57], our finding that CD4^+^ T cells are vital for the recruitment of CD8^+^ T cells to the tumor site is perhaps not unexpected. Surprisingly, we observed fewer tumor-infiltrating CD4^+^ T cells in mice lacking CD8^+^ T cells. It is unclear how CD8^+^ T cells might affect the trafficking of CD4^+^ T cells to the tumor site. Additional studies are necessary to address the trafficking and function of CD4^+^ T cells in the immune response to αKO KPC tumors.

Although PIK3CG is expressed mainly in hematopoietic cells, it was reported to be present at high levels in human pancreatic cancer tissue. Downregulation of PIK3CG slightly slowed the proliferation of PDAC cell lines in culture [58]. A subsequent study showed that genetic ablation or systemic pharmacological inhibition of PIK3CG led to activation of T cells and slower PDAC growth *in vivo* [21]. These antitumor effects were attributed to suppression of PIK3CG in macrophages leading to suppression of cytotoxic T cells and not due to decreased PIK3CG in tumor cells. Genetic or pharmacologic downregulation of PIK3CG altered the transcriptional program of tumor-infiltrating macrophages to promote cytotoxic T cell infiltration of pancreatic tumors, thus extending survival of the mice [21, 59, 60]. However, unlike ablation of *Pik3ca* in KPC tumor cells, all of the animals with downregulated PIK3CG eventually died from pancreatic cancer progression [21]. PIK3CD is expressed mainly in leukocytes and is critical for the differentiation and function of effector T cells and CD4^+^ regulatory T cells (Tregs) [61]. Suppression of PIK3CD in Tregs protected host mice from a broad range of transplanted cancers [20], but it is unclear if downregulation of PIK3CD slowed pancreatic cancer growth *in vivo* [62]. Several immunotherapeutic strategies are being investigated to promote infiltration of T cells into pancreatic tumors [10]. Our results indicate that PIK3CA may be a potential drug target to increase T cell immunogenicity of pancreatic cancer. Treatment with a pan-isoform PI3K inhibitor did not reduce the tumor volume of KPC cells or patient-derived xenografts implanted in mice [63]. Based on our current understanding, systemic pharmacological inhibition of all PI3K isoforms should enhance the immunogenicity of the pancreatic tumors by upregulating MHC I and CD80, but inhibitory effects on cytotoxic T cells will negate these beneficial changes.

In conclusion, we have shown that PIK3CA-AKT signaling in WT KPC tumors reduces the cell surface expression of MHC I and CD80 to promote immune evasion. Genetic ablation of *Pik3ca* counteracts these effects to render pancreatic tumors more immunogenic and thus more susceptible to T cell clearance. Some tumor cells have been demonstrated to directly present tumor antigens to CD8^+^ T cells [64, 65]. Whether αKO cells directly present tumor antigens on their surface is still unclear. In our current model, we believe that abrogating PIK3CA-AKT signaling in tumor cells attracts both CD4^+^ and CD8^+^ T cells to infiltrate the tumors. CD4^+^ and CD8^+^ T cells act synergistically to cause tumor regression. WT KPC tumors are surrounded by T cells and evade the immune system due to PIK3CA-mediated downregulation of MHC I and CD80. αKO tumors, on the other hand, are infiltrated with T cells that can recognize and lyse the tumors. Our results could pave the way for a robustly effective treatment for pancreatic cancer, which has so far been lacking.

## MATERIALS AND METHODS

### Cell Lines

FC1245 cells [25] were a gift from Dr. David Tuveson (Cold Spring Harbor Laboratory). SNP analysis showed that the cells have a 99.63% C57BL/6J and 0.37% C57BL/6NJ genetic background with a single T/A polymorphism on chromosome 8. FC1245 cells were infected with lentiviral particles containing firefly luciferase under the control of a CMV promoter (Cellomics Technology, PLV-10064). 48 hours after infection, cells were selected with 1.5 mg/mL G418 and screened for luciferase expression on the IVIS Lumina III imaging system (Xenogen) to generate stable luciferase-expressing cells referred to as wildtype (WT) KPC cells. To generate *Pik3ca*^-/-^ (αKO) KPC cell lines, WT KPC cells were transfected concurrently with *Pik3ca* CRISPR/Cas9 KO and HDR plasmids (Santa Cruz Biotechnology, sc-422231 and sc-422231-HDR). To generate the *Egfr*^/^ (EgfrKO) cell line, WT KPC cells were transfected concurrently with *Egfr* CRISPR/Cas9 KO and HDR plasmids (Santa Cruz Biotechnology, sc-420131 and sc-420131-HDR). Transfected cells were selected with 5 μg/mL puromycin and RFP^+^ cells were collected using fluorescence-activated cell sorting (FACS) on a FACSAria (BD Biosciences). RFP^+^ cells were serially diluted in a 96-well plate to generate single cell clones. Clones that showed an absence of PIK3CA or EGFR protein on western blots were retained. αKO KPC cells lacking CD80 (referred to as αKO/CD80KO) were generated by transfecting αKO cells with B7-1 CRISPR/Cas9 KO plasmid (Santa Cruz Biotechnology, sc-419570). 48 hours after transfection, cells were collected using StemPro Accutase (ThermoFisher Scientific), incubated with FcR block, and stained for CD80. CD80-null cells were collected by FACS. Single cell clones were generated by serial dilution and confirmed by flow cytometry. αKO cells with reduced levels of B2m (referred to as αKO/shB2m) were generated by infecting αKO cells with B2m shRNA lentiviral particles (Santa Cruz Biotechnology, sc-29593-V). 48 hours after infection, the cells were stained for H-2K^b^ and cells with low levels of H-2K^b^ were collected by FACS. αKO/CD80KO cells were infected with β-2-microglobulin shRNA lentiviruses and FACS sorted to generate αKO KPC cells without CD80 and with low levels of H-2K^b^ (referred to as αKO/CD80KO+shB2m). αKO-caAkt and αKO-Cont cells were generated by infecting αKO cells with lentiviral particles expressing a constitutively active human Akt1 mutant and EGFP or EGFP alone, respectively (pHRIG-Akt1 (Addgene plasmid #53583) was a gift from Heng Zhao [66]). 48 hours after infection, cells expressing EGFP were collected using FACS; mutant Akt1 expression was confirmed by western blotting. All murine cell lines were cultured in DMEM media containing 10% fetal bovine serum and 1% penicillin/streptomycin at 37 °C and 5% CO_2_. Human pancreatic cancer cell lines Panc1, HPAF-II, and PSN1 were purchased from ATCC and cultured using recommended conditions. Patient-derived human pancreatic cancer cell lines were previously described and cultured as recommended [67].

### 2D Cell Proliferation Assay

5,000 cells were plated in triplicate in a 6-well plate. On days 1 through 4, cells were trypsinized, mixed with an equal volume of trypan blue, and counted using the Countess II automated cell counter (Life Technologies). Cell number was plotted against time using GraphPad Prism 7.

### 3D Cell Culture

3D cultures used previously described protocols [68]. Briefly, 1,000 cells in DMEM containing 10% FBS were mixed with methylcellulose stock solution to a final concentration of 0.24% methylcellulose. Cells were cultured by the hanging drop method on the lid of a 96-well plate. To prevent evaporation, PBS was added to the plate. The plates were incubated at 37 °C in a 5% CO_2_ atmosphere. The cells were imaged on day 5 using brightfield microscopy with a 40X objective.

### Immunoblotting

Cell lysates were prepared in RIPA buffer (50 mM HEPES, pH 7.4, 10 mM sodium pyrophosphate, 50 mM NaCl, 50 mM NaF, 5 mM EDTA, 1 mM sodium orthovanadate, 0.25% sodium deoxycholate, 1% NP40, 1 mM PMSF, and protease inhibitor cocktail (Sigma P8340)) and cleared by centrifugation at 4 °C. Cell proteins were separated by SDS-PAGE and transferred to nitrocellulose membranes by semi-dry transfer, and the membranes were incubated with primary antibodies in Tris-buffered saline plus 0.1% Tween 20. After incubation in HRP-linked secondary antibody followed by ECL reagent (Western Lightning Plus-ECL (PerkinElmer) or SuperSignal West Femto (ThermoFisher Scientific), signals were detected using a FluorChem E imager (ProteinSimple). To detect multiple proteins on the same membrane, HRP was inactivated by incubating membranes in 30% H_2_O_2_ for 30 minutes [69], or the membranes were stripped for 30 min at 50 °C in 62.5 mM Tris, pH 6.7, 2% SDS, and 100 mM 2-mercaptoethanol prior to reprobing with another antibody.

### Reverse Phase Protein Array (RPPA)

Cells were plated in duplicate in a 6-well plate, washed 2 times in PBS, and lysed in RPPA lysis buffer (1% Triton X-100, 50 mM HEPES, pH 7.4, 150 mM NaCl, 1.5 mM MgCl_2_, 1 mM EGTA, 100 mM NaF, 10 mM sodium pyrophosphate, 1 mM sodium orthovanadate, 10% glycerol, 1 mM PMSF, and protease inhibitor cocktail). Protein concentration was adjusted to 1.5 μg/μl in SDS sample buffer (40% glycerol, 8% SDS, 0.25 M Tris-HCl, pH 6.8, and 1.43 M 2-mercaptoethanol). The samples were sent to M.D. Anderson Cancer Center Core Facility for RPPA analysis. Briefly, cell lysates were serially diluted two-fold for 5 dilutions (from undiluted to 1:16 dilution) and arrayed on nitrocellulose-coated slides in an 11×11 format. Samples were probed with antibodies using a tyramide-based signal amplification approach and visualized by DAB colorimetric reaction. Slides were scanned on a flatbed scanner to produce 16-bit tiff images. Spots from tiff images were identified and the density was quantified by Array-Pro Analyzer. Normalized Log2 values were median-centered and used for heatmap generation using GraphPad Prism v7.

### Mice

C57BL/6J (B6; stock# 000664), B6.CB17-*Prkdc^scid^*/SzJ (SCID; stock# 001913), B6.129S2-*Cd4^tm1Mak^*/J (CD4KO; stock# 002663), and B6.129S2-*Cd8a^tm1Mak^*/J (CD8KO; stock# 002665) mice were purchased from Jackson Laboratories. All animals are in the B6 genetic background. Mice used for orthotopic implantation and adoptive transfer recipients were males 8-10 weeks of age. Experiments were conducted in accordance with the Office of Laboratory Animal Welfare and approved by the Institutional Animal Care and Use Committee of Stony Brook University.

### Orthotopic Implantation and Tumor Imaging

Cells were trypsinized and washed twice in PBS. Mice were anesthetized with a mixture of 100 mg/kg ketamine and 10 mg/kg xylazine. The abdomen was shaved and swabbed with a sterile alcohol pad followed by povidone-iodide scrub. A small vertical incision was made over the left lateral abdominal area, to the left of the spleen. The head of the pancreas attached to the duodenum was located. Using a sterile Hamilton syringe with a 27 gauge needle, 0.5 million cells in 30 μl PBS were injected into the head of the pancreas. The injection site was pressed with a sterile cotton swab to prevent leakage. The abdominal and skin incisions were closed with 5-0 silk black braided sutures. The mice were given an intraperitoneal injection of 2 mg/kg Ketorolac immediately after surgery. To monitor tumor growth, the animals were injected intraperitoneally with 100 mg/kg RediJect D-Luciferin (PerkinElmer) and imaged on the IVIS Lumina III imaging system (Xenogen). Data were analyzed using Living Image v4.3.1 software.

### Survival Studies

Mice implanted with tumor cells were monitored by IVIS imaging every week for tumor growth. Death, weight loss of ≥ 15% body weight, or inability to move were considered as endpoints at which surviving animals were euthanized.

### Neutralizing Antibody Experiment

B6 mice were injected intraperitoneally with 500 μg of neutralizing antibodies against CD4 (BioXcell, Clone GK1.5) and CD8 (BioXcell, Clone 53-5.8) antigens on Days 1 and 3. On day 3, depletion of CD4 and CD8 T cells was confirmed by flow cytometry. The control mouse received PBS injections. On day 4, mice were implanted with 0.5 million αKO cells in the head of the pancreas as before. Following implantation, mice received weekly injections of neutralizing antibodies till the end of the study. Tumor growth was monitored by IVIS imaging.

### Adoptive T Cell Transfer

Six months after B6 mice were implanted with αKO cells, the mice were euthanized and their spleens collected under sterile conditions in PBS. The spleen was forced through a 70 μm filter and washed with 10 ml PBS. RBCs were lysed using ACK lysis buffer (150 mM NH_4_Cl, 10 mM KHCO_3_, and 0.1 mM Na_2_EDTA, pH 7.2-7.4) and the cells were washed 2 times in PBS and collected by centrifugation at 330 × g for 5 minutes at 4 °C. Splenocytes were counted and resuspended in MACS buffer for T cell isolation using the Mouse Pan T Cell Isolation Kit II (Miltenyi, #130-095-130) following the manufacturer’s instructions. Cells were analyzed for CD90.2 expression by flow cytometry after purification to ensure ≥98% purity. Purified T cells from 2 or more mice were pooled and the desired number of cells was suspended in 100 μl PBS and injected retro-orbitally into SCID mice.

### Flow Cytometry

Cells were detached from the tissue culture plate using trypsin-EDTA (Corning) or StemPro Accutase (CD80 and H-2K^b^ staining). They were washed 2 times in PBS and incubated with 1:50 FcR block for 20 minutes prior to staining for 30 minutes with antibodies. Cells were washed twice in PBS and immediately analyzed by flow cytometry on a FACScalibur (BD Biosciences) or DxP 8 (Cytek). Cells to be analyzed at a later time were fixed in 2% paraformaldehyde. Data were analyzed using FlowJo.

### Histology

Mouse organs were fixed in 10% formalin for 24 hours. Tissues were processed using a Leica ASP300S Tissue Processor, paraffin embedded, and cut into 4-μm sections. H&E staining was performed using Hematoxylin Solution, Gill No.3 (Sigma-Aldrich) and Eosin Y (Fisher Scientific). Immunohistochemistry (IHC) was performed manually following deparaffinization and rehydration. Antigen retrieval was done in citrate buffer pH 6.0 (Vector Laboratories) using a Decloaking Chamber (Biocare Medical). Endogenous peroxidase activity and biotin were blocked using H_2_O_2_ and the Avidin/Biotin Blocking Kit (Vector Laboratories), respectively. Following incubation with primary antibodies, the R.T.U. Vectastain Kit (Vector Laboratories) and 3,3’-diaminobenzidine in chromogen solution (Dako) were used to develop the signal. The sections were then counterstained with hematoxylin (Dako). For quantification of T cells, ImageJ was used to count cells in eight 20X microscopic fields per mouse.

### Antibodies

*Western blotting:* PIK3CA (#4249S), pAKT T308 (#9275S), pAKT S473(#4051S), and β-actin (#4970S) were from Cell Signaling Technology. AKT (Santa Cruz, sc-8312), β-tubulin (Sigma-Aldrich, T4026), and EGFR (Epitomics, 1902-1). Anti-mouse IgG secondary antibody, HRP (62-6520) and anti-rabbit IgG secondary antibody, HRP (31460) were from ThermoFisher Scientific. *Flow cytometry:* anti-mouse FcR block (#101302), CD90.2 (clone 30-H12), anti-mouse CD80 (clone 16-10A1), H-2K^b^ (clone AF6-88.5), Human Trustain FcX (#422301), and anti-human CD80 (clone 2D10) were from BioLegend. HLA-ABC (eBioscience, Clone W6/32), annexin V (Invitrogen #A23204). *Histology:* F4/80 (ab111101), CD3 (ab16669) and CD4 (ab183685) were from Abcam. CD8 (Cell Signaling Technology, #98941).

### Inhibitor Studies

200,000 KPC cells or 0.5 million human pancreatic cancer cells were plated per well in a 6-well plate. 10 μM Akt inhibitor VIII (EMD Millipore, 124017) was added to cells the following day in fresh media. For the dose-response curve, cells were treated with increasing concentrations of Akt inhibitor or DMSO. Cells were incubated at 37 °C and 5% CO_2_ for 48 hours. Cells were detached from the plate using StemPro Accutase, washed 2 times in PBS, incubated with FcR block, stained with anti-CD80 and anti-H-2K^b^ (mouse) or anti-HLA-ABC and anti-CD80 (human) antibodies, and analyzed by flow cytometry (FACScalibur or DxP 8). Cells from a duplicate plate were lysed and subjected to western blotting.

### Quantitative Real-time PCR (qRT-PCR)

RNA was extracted from cells using the RNeasy Kit (Qiagen). The RNase-free DNase Set (Qiagen) was used for on-column DNase digestion. cDNA was synthesized using the iScript cDNA Synthesis Kit (Bio-Rad). A StepOnePlus Real-Time PCR system was used for qRT-PCR. All reactions were carried out in triplicate using TaqMan gene expression assays. *Hprt* (Mm00446068_m1), *B2m* (Mm00437762_m1), *CD80* (Mm00711660_m1), and *H-2K^b^* (Mm01612247_mH2-K1) were purchased from ThermoFisher Scientific. Gene expression changes were normalized to *Hprt* and analyzed by the 2^-ΔCT^ method.

### TCGA Analysis

The TCGA data were downloaded as z-scores from the cBioPortal (http://www.cbioportal.org). Patient samples were rank ordered based on their PTEN or pAKT S473 expression. Samples with high and low mRNA/protein expression were used for comparing the levels of CD80, B2m, or HLA-ABC.

### Statistics

All statistical analyses were done using GraphPad Prism v7.

## AUTHOR CONTRIBUTIONS

N.S designed and carried out experiments, analyzed and interpreted the data, wrote the manuscript with help from A.W.M.V, R.Z.L, and L.M.B.

P.A.M designed experiments, helped with adoptive transfers and flow cytometry experiments and analysis.

H.V.H helped with *in vitro* inhibitor experiments.

O.P performed TCGA analysis.

Y.J performed all animal surgeries.

L.M.B helped design experiments, analyze and interpret data, and writing of the manuscript.

K.P and C.L provided human pancreatic cancer cell lines, helped with data interpretation.

A.W.M.V helped with experimental design, analysis, interpretation of data, and manuscript writing.

R.Z.L supervised the project, designed experiments, analyzed and interpreted data, and helped writing the manuscript.

## ACKNOWLEDGEMENTS

This work was funded by NIH grants DK108989 (RZL), CA194836 (RZL), and AI101221 (AVDV) and a VA Merit Award BX004083 (RZL). Reverse Phase Protein Array (RPPA) was performed by the RPPA core facility at MD Anderson Cancer center, funded by NCI grant CA16672. We thank Juei-Suei Chen for help with H&E staining and immunohistochemistry.

## References

1. NCI. https://seer.cancer.gov/statfacts/html/pancreas.html

2. Hackert, T., M.W. Buchler, and J. Werner, Surgical options in the management of pancreatic cancer. Minerva Chir, 2009. 64(5): p. 465–76.

3. Balachandran, V.P., M. Luksza, J.N. Zhao, V. Makarov, J.A. Moral, R. Remark, et al., Identification of unique neoantigen qualities in long-term survivors of pancreatic cancer. Nature, 2017. 551(7681): p. 512–516.

4. Fukunaga, A., M. Miyamoto, Y. Cho, S. Murakami, Y. Kawarada, T. Oshikiri, et al., CD8+ tumor-infiltrating lymphocytes together with CD4+ tumor-infiltrating lymphocytes and dendritic cells improve the prognosis of patients with pancreatic adenocarcinoma. Pancreas, 2004. 28(1): p. e26–31.

5. Ino, Y., R. Yamazaki-Itoh, K. Shimada, M. Iwasaki, T. Kosuge, Y. Kanai, et al., Immune cell infiltration as an indicator of the immune microenvironment of pancreatic cancer. Br J Cancer, 2013. 108(4): p. 914–23.

6. Lohneis, P., M. Sinn, S. Bischoff, A. Juhling, U. Pelzer, L. Wislocka, et al., Cytotoxic tumour-infiltrating T lymphocytes influence outcome in resected pancreatic ductal adenocarcinoma. Eur J Cancer, 2017. 83: p. 290–301.

7. Ekkirala, C.R., P. Cappello, R.S. Accolla, M. Giovarelli, I. Romero, C. Garrido, et al., Class II transactivator-induced MHC class II expression in pancreatic cancer cells leads to tumor rejection and a specific antitumor memory response. Pancreas, 2014. 43(7): p. 1066–72.

8. Ryschich, E., T. Notzel, U. Hinz, F. Autschbach, J. Ferguson, I. Simon, et al., Control of T-cell-mediated immune response by HLA class I in human pancreatic carcinoma. Clin Cancer Res, 2005. 11(2 Pt 1): p. 498–504.

9. Pandha, H., A. Rigg, J. John, and N. Lemoine, Loss of expression of antigen-presenting molecules in human pancreatic cancer and pancreatic cancer cell lines. Clin Exp Immunol, 2007. 148(1): p. 127–35.

10. Bengsch, F., D.M. Knoblock, A. Liu, F. McAllister, and G.L. Beatty, CTLA-4/CD80 pathway regulates T cell infiltration into pancreatic cancer. Cancer Immunol Immunother, 2017. 66(12): p. 1609–1617.

11. Putzer, B.M., F. Rodicker, M.M. Hitt, T. Stiewe, and H. Esche, Improved treatment of pancreatic cancer by IL-12 and B7.1 costimulation: antitumor efficacy and immunoregulation in a nonimmunogenic tumor model. Mol Ther, 2002. 5(4): p. 405–12.

12. Thomas, G.R. and J. Wen, Endogenous expression of CD80 co-stimulatory molecule facilitates in vivo tumor regression of oral squamous carcinoma. Anticancer Res, 2006. 26(6B): p. 4093–101.

13. Hruban, R.H., C. Iacobuzio-Donahue, R. E. Wilentz, M. Goggins, and S.E. Kern, Molecular pathology of pancreatic cancer. Cancer J, 2001. 7(4): p. 251–8.

14. Janku, F., Phosphoinositide 3-kinase (PI3K) pathway inhibitors in solid tumors: From laboratory to patients. Cancer Treat Rev, 2017. 59: p. 93–101.

15. Engelman, J.A., Targeting PI3K signalling in cancer: opportunities, challenges and limitations. Nat Rev Cancer, 2009. 9(8): p. 550–62.

16. Fruman, D.A. and C. Rommel, PI3K and cancer: lessons, challenges and opportunities. Nat Rev Drug Discov, 2014. 13(2): p. 140–56.

17. Murthy, D., K.S. Attri, and P.K. Singh, Phosphoinositide 3-Kinase Signaling Pathway in Pancreatic Ductal Adenocarcinoma Progression, Pathogenesis, and Therapeutics. Front Physiol, 2018. 9: p. 335.

18. Wu, C.Y., E.S. Carpenter, K.K. Takeuchi, C.J. Halbrook, L.V. Peverley, H. Bien, et al., PI3K regulation of RAC1 is required for KRAS-induced pancreatic tumorigenesis in mice. Gastroenterology, 2014. 147(6): p. 1405–16 e7.

19. Baer, R., C. Cintas, M. Dufresne, S. Cassant-Sourdy, N. Schonhuber, L. Planque, et al., Pancreatic cell plasticity and cancer initiation induced by oncogenic Kras is completely dependent on wild-type PI 3-kinase p110alpha. Genes Dev, 2014. 28(23): p. 2621–35.

20. Ali, K., D.R. Soond, R. Pineiro, T. Hagemann, W. Pearce, E.L. Lim, et al., Inactivation of PI(3)K p110delta breaks regulatory T-cell-mediated immune tolerance to cancer. Nature, 2014. 510(7505): p. 407–411.

21. Kaneda, M.M., P. Cappello, A.V. Nguyen, N. Ralainirina, C.R. Hardamon, P. Foubert, et al., Macrophage PI3Kgamma Drives Pancreatic Ductal Adenocarcinoma Progression. Cancer Discov, 2016. 6(8): p. 870–85.

22. Baer, R., C. Cintas, N. Therville, and J. Guillermet-Guibert, Implication of PI3K/Akt pathway in pancreatic cancer: When PI3K isoforms matter? Adv Biol Regul, 2015. 59: p. 19–35.

23. Alagesan, B., G. Contino, A.R. Guimaraes, R.B. Corcoran, V. Deshpande, G.R. Wojtkiewicz, et al., Combined MEK and PI3K inhibition in a mouse model of pancreatic cancer. Clin Cancer Res, 2015. 21(2): p. 396–404.

24. Hingorani, S.R., L. Wang, A.S. Multani, C. Combs, T.B. Deramaudt, R.H. Hruban, et al., Trp53R172H and KrasG12D cooperate to promote chromosomal instability and widely metastatic pancreatic ductal adenocarcinoma in mice. Cancer Cell, 2005. 7(5): p. 469–83.

25. Roy, I., D.M. McAllister, E. Gorse, K. Dixon, C.T. Piper, N.P. Zimmerman, et al., Pancreatic Cancer Cell Migration and Metastasis Is Regulated by Chemokine-Biased Agonism and Bioenergetic Signaling. Cancer Res, 2015. 75(17): p. 3529–42.

26. Ardito, C.M., B.M. Gruner, K.K. Takeuchi, C. Lubeseder-Martellato, N. Teichmann, P.K. Mazur, et al., EGF receptor is required for KRAS-induced pancreatic tumorigenesis. Cancer Cell, 2012. 22(3): p. 304–17.

27. Chung, W.J., A. Daemen, J.H. Cheng, J.E. Long, J.E. Cooper, B.E. Wang, et al., Kras mutant genetically engineered mouse models of human cancers are genomically heterogeneous. Proc Natl Acad Sci U S A, 2017. 114(51): p. E10947–E10955.

28. Qiu, W., F. Sahin, C.A. Iacobuzio-Donahue, D. Garcia-Carracedo, W.M. Wang, C.Y. Kuo, et al., Disruption of p16 and activation of Kras in pancreas increase ductal adenocarcinoma formation and metastasis in vivo. Oncotarget, 2011. 2(11): p. 862–73.

29. Uchiyama, M., N. Usami, M. Kondo, S. Mori, M. Ito, G. Ito, et al., Loss of heterozygosity of chromosome 12p does not correlate with KRAS mutation in non-small cell lung cancer. Int J Cancer, 2003. 107(6): p. 962–9.

30. Yu, C.C., W. Qiu, C.S. Juang, M.M. Mansukhani, B. Halmos, and G.H. Su, Mutant allele specific imbalance in oncogenes with copy number alterations: Occurrence, mechanisms, and potential clinical implications. Cancer Lett, 2017. 384: p. 86–93.

31. Burgess, M.R., E. Hwang, R. Mroue, C.M. Bielski, A.M. Wandler, B.J. Huang, et al., KRAS Allelic Imbalance Enhances Fitness and Modulates MAP Kinase Dependence in Cancer. Cell, 2017. 168(5): p. 817–829 e15.

32. Marzo, A.L., B.F. Kinnear, R.A. Lake, J.J. Frelinger, E.J. Collins, B.W. Robinson, et al., Tumor-specific CD4+ T cells have a major “post-licensing” role in CTL mediated anti-tumor immunity. J Immunol, 2000. 165(11): p. 6047–55.

33. Durrant, L.G., K.C. Ballantyne, N.C. Armitage, R.A. Robins, R. Marksman, J.D. Hardcastle, et al., Quantitation of MHC antigen expression on colorectal tumours and its association with tumour progression. Br J Cancer, 1987. 56(4): p. 425–32.

34. Tabibzadeh, S.S., A. Sivarajah, D. Carpenter, B.M. Ohlsson-Wilhelm, and P.G. Satyaswaroop, Modulation of HLA-DR expression in epithelial cells by interleukin 1 and estradiol-17 beta. J Clin Endocrinol Metab, 1990. 71(3): p. 740–7.

35. Clark, C.E., S.R. Hingorani, R. Mick, C. Combs, D.A. Tuveson, and R.H. Vonderheide, Dynamics of the immune reaction to pancreatic cancer from inception to invasion. Cancer Res, 2007. 67(19): p. 9518–27.

36. Ene-Obong, A., A.J. Clear, J. Watt, J. Wang, R. Fatah, J.C. Riches, et al., Activated Pancreatic Stellate Cells Sequester CD8(+) T-Cells to Reduce Their Infiltration of the Juxtatumoral Compartment of Pancreatic Ductal Adenocarcinoma. Gastroenterology, 2013. 145(5): p. 1121–1132.

37. Shibuya, K.C., V.K. Goel, W. Xiong, J.G. Sham, S.M. Pollack, A.M. Leahy, et al., Pancreatic ductal adenocarcinoma contains an effector and regulatory immune cell infiltrate that is altered by multimodal neoadjuvant treatment. PLoS One, 2014. 9(5): p. e96565.

38. Beatty, G.L., E.G. Chiorean, M.P. Fishman, B. Saboury, U.R. Teitelbaum, W. Sun, et al., CD40 agonists alter tumor stroma and show efficacy against pancreatic carcinoma in mice and humans. Science, 2011. 331(6024): p. 1612–6.

39. Aiello, N.M., D.L. Bajor, R.J. Norgard, A. Sahmoud, N. Bhagwat, M.N. Pham, et al., Metastatic progression is associated with dynamic changes in the local microenvironment. Nat Commun, 2016. 7: p. 12819.

40. Park, I.A., S.H. Hwang, I.H. Song, S.H. Heo, Y.A. Kim, W.S. Bang, et al., Expression of the MHC class II in triple-negative breast cancer is associated with tumor-infiltrating lymphocytes and interferon signaling. PLoS One, 2017. 12(8): p. e0182786.

41. Brea, E.J., C.Y. Oh, E. Manchado, S. Budhu, R.S. Gejman, G. Mo, et al., Kinase Regulation of Human MHC Class I Molecule Expression on Cancer Cells. Cancer Immunol Res, 2016. 4(11): p. 936–947.

42. Zhou, F., Molecular mechanisms of IFN-gamma to up-regulate MHC class I antigen processing and presentation. Int Rev Immunol, 2009. 28(3-4): p. 239–60.

43. Swann, S.A., M. Williams, C.M. Story, K.R. Bobbitt, R. Fleis, and K.L. Collins, HIV-1 Nef blocks transport of MHC class I molecules to the cell surface via a PI 3-kinase-dependent pathway. Virology, 2001. 282(2): p. 267–77.

44. Allison, J.P., A.A. Hurwitz, and D.R. Leach, Manipulation of costimulatory signals to enhance antitumor T-cell responses. Curr Opin Immunol, 1995. 7(5): p. 682–6.

45. Tirapu, I., E. Huarte, C. Guiducci, A. Arina, M. Zaratiegui, O. Murillo, et al., Low surface expression of B7-1 (CD80) is an immunoescape mechanism of colon carcinoma. Cancer Res, 2006. 66(4): p. 2442–50.

46. Jiang, Y., Y. Li, and B. Zhu, T-cell exhaustion in the tumor microenvironment. Cell Death Dis, 2015. 6: p. e1792.

47. Sun, C., R. Mezzadra, and T.N. Schumacher, Regulation and Function of the PD-L1 Checkpoint. Immunity, 2018. 48(3): p. 434–452.

48. Xing, X., J. Guo, X. Wen, G. Ding, B. Li, B. Dong, et al., Analysis of PD1, PDL1, PDL2 expression and T cells infiltration in 1014 gastric cancer patients. Oncoimmunology, 2018. 7(3): p. e1356144.

49. Bos, R. and L.A. Sherman, CD4+ T-cell help in the tumor milieu is required for recruitment and cytolytic function of CD8+ T lymphocytes. Cancer Res, 2010. 70(21): p. 8368–77.

50. Dranoff, G., E. Jaffee, A. Lazenby, P. Golumbek, H. Levitsky, K. Brose, et al., Vaccination with irradiated tumor cells engineered to secrete murine granulocyte-macrophage colony-stimulating factor stimulates potent, specific, and long-lasting anti-tumor immunity. Proc Natl Acad Sci U S A, 1993. 90(8): p. 3539–43.

51. Tassi, E., F. Gavazzi, L. Albarello, V. Senyukov, R. Longhi, P. Dellabona, et al., Carcinoembryonic antigen-specific but not antiviral CD4+ T cell immunity is impaired in pancreatic carcinoma patients. J Immunol, 2008. 181(9): p. 6595–603.

52. Quezada, S.A., T.R. Simpson, K.S. Peggs, T. Merghoub, J. Vider, X. Fan, et al., Tumor-reactive CD4(+) T cells develop cytotoxic activity and eradicate large established melanoma after transfer into lymphopenic hosts. J Exp Med, 2010. 207(3): p. 637–50.

53. Xie, Y., A. Akpinarli, C. Maris, E.L. Hipkiss, M. Lane, E.K. Kwon, et al., Naive tumor-specific CD4(+) T cells differentiated in vivo eradicate established melanoma. J Exp Med, 2010. 207(3): p. 651–67.

54. Janssen, E.M., E.E. Lemmens, T. Wolfe, U. Christen, M.G. von Herrath, and S.P. Schoenberger, CD4+ T cells are required for secondary expansion and memory in CD8+ T lymphocytes. Nature, 2003. 421(6925): p. 852–6.

55. Shedlock, D.J. and H. Shen, Requirement for CD4 T cell help in generating functional CD8 T cell memory. Science, 2003. 300(5617): p. 337–9.

56. Li, K., M. Baird, J. Yang, C. Jackson, F. Ronchese, and S. Young, Conditions for the generation of cytotoxic CD4(+) Th cells that enhance CD8(+) CTL-mediated tumor regression. Clin Transl Immunology, 2016. 5(8): p. e95.

57. Wong, S.B., R. Bos, and L.A. Sherman, Tumor-specific CD4+ T cells render the tumor environment permissive for infiltration by low-avidity CD8+ T cells. J Immunol, 2008. 180(5): p. 3122–31.

58. Edling, C.E., F. Selvaggi, R. Buus, T. Maffucci, P. Di Sebastiano, H. Friess, et al., Key role of phosphoinositide 3-kinase class IB in pancreatic cancer. Clin Cancer Res, 2010. 16(20): p. 4928–37.

59. Kaneda, M.M., K.S. Messer, N. Ralainirina, H. Li, C.J. Leem, S. Gorjestani, et al., PI3Kgamma is a molecular switch that controls immune suppression. Nature, 2016. 539(7629): p. 437–442.

60. Kaneda, M.M., K.S. Messer, N. Ralainirina, H. Li, C.J. Leem, S. Gorjestani, et al., Corrigendum: PI3Kgamma is a molecular switch that controls immune suppression. Nature, 2017. 542(7639): p. 124.

61. Stark, A.K., S. Sriskantharajah, E.M. Hessel, and K. Okkenhaug, PI3K inhibitors in inflammation, autoimmunity and cancer. Curr Opin Pharmacol, 2015. 23: p. 82–91.

62. Ali, K., D.R. Soond, R. Pineiro, T. Hagemann, W. Pearce, E.L. Lim, et al., Corrigendum: Inactivation of PI(3)K p110delta breaks regulatory T-cell-mediated immune tolerance to cancer. Nature, 2016. 535(7613): p. 580.

63. Eser, S., N. Reiff, M. Messer, B. Seidler, K. Gottschalk, M. Dobler, et al., Selective requirement of PI3K/PDK1 signaling for Kras oncogene-driven pancreatic cell plasticity and cancer. Cancer Cell, 2013. 23(3): p. 406–20.

64. Hargadon, K.M., C.C. Brinkman, L. Sheasley-O’neill S, L.A. Nichols, T.N. Bullock, and V.H. Engelhard, Incomplete differentiation of antigen-specific CD8 T cells in tumor-draining lymph nodes. J Immunol, 2006. 177(9): p. 6081–90.

65. Wolkers, M.C., G. Stoetter, F.A. Vyth-Dreese, and T.N. Schumacher, Redundancy of direct priming and cross-priming in tumor-specific CD8+ T cell responses. J Immunol, 2001. 167(7): p. 3577–84.

66. Xie, R., M. Cheng, M. Li, X. Xiong, M. Daadi, R.M. Sapolsky, et al., Akt isoforms differentially protect against stroke-induced neuronal injury by regulating mTOR activities. J Cereb Blood Flow Metab, 2013. 33(12): p. 1875–85.

67. Pham, K., D. Delitto, A.E. Knowlton, E.R. Hartlage, R. Madhavan, D.H. Gonzalo, et al., Isolation of Pancreatic Cancer Cells from a Patient-Derived Xenograft Model Allows for Practical Expansion and Preserved Heterogeneity in Culture. Am J Pathol, 2016. 186(6): p. 1537–46.

68. Longati, P., X. Jia, J. Eimer, A. Wagman, M.R. Witt, S. Rehnmark, et al., 3D pancreatic carcinoma spheroids induce a matrix-rich, chemoresistant phenotype offering a better model for drug testing. BMC Cancer, 2013. 13: p. 95.

69. Sennepin, A.D., S. Charpentier, T. Normand, C. Sarre, A. Legrand, and L.M. Mollet, Multiple reprobing of Western blots after inactivation of peroxidase activity by its substrate, hydrogen peroxide. Anal Biochem, 2009. 393(1): p. 129–31.

