## Supplemental Information for "PIK3CA in *Kras^G12D^/Trp53^R172H^* Tumor Cells Promotes Immune Evasion by Limiting Infiltration of T Cells in a Model of Pancreatic Cancer"

### Supplementary Information

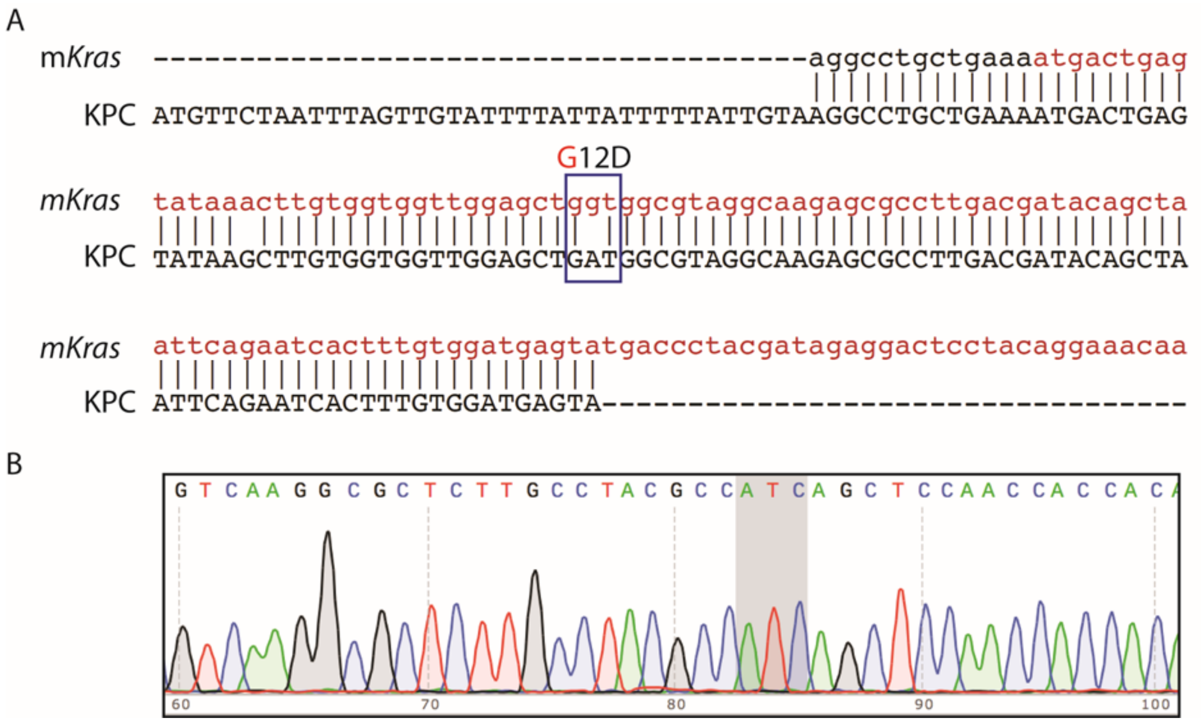


**Supplementary Fig. S1. Sequencing of *Kras* Exon1.** Genomic DNA from WT KPC cells was used for sequencing exon 1 of the *Kras* gene. **(A)** Sequence alignment of DNA from KPC cells (KPC) against murine *Kras* gene (NCBI Gene ID 16653) (m*Kras*). **(B)** Chromatogram showing G12D mutation (grey highlight) in WT KPC cells.

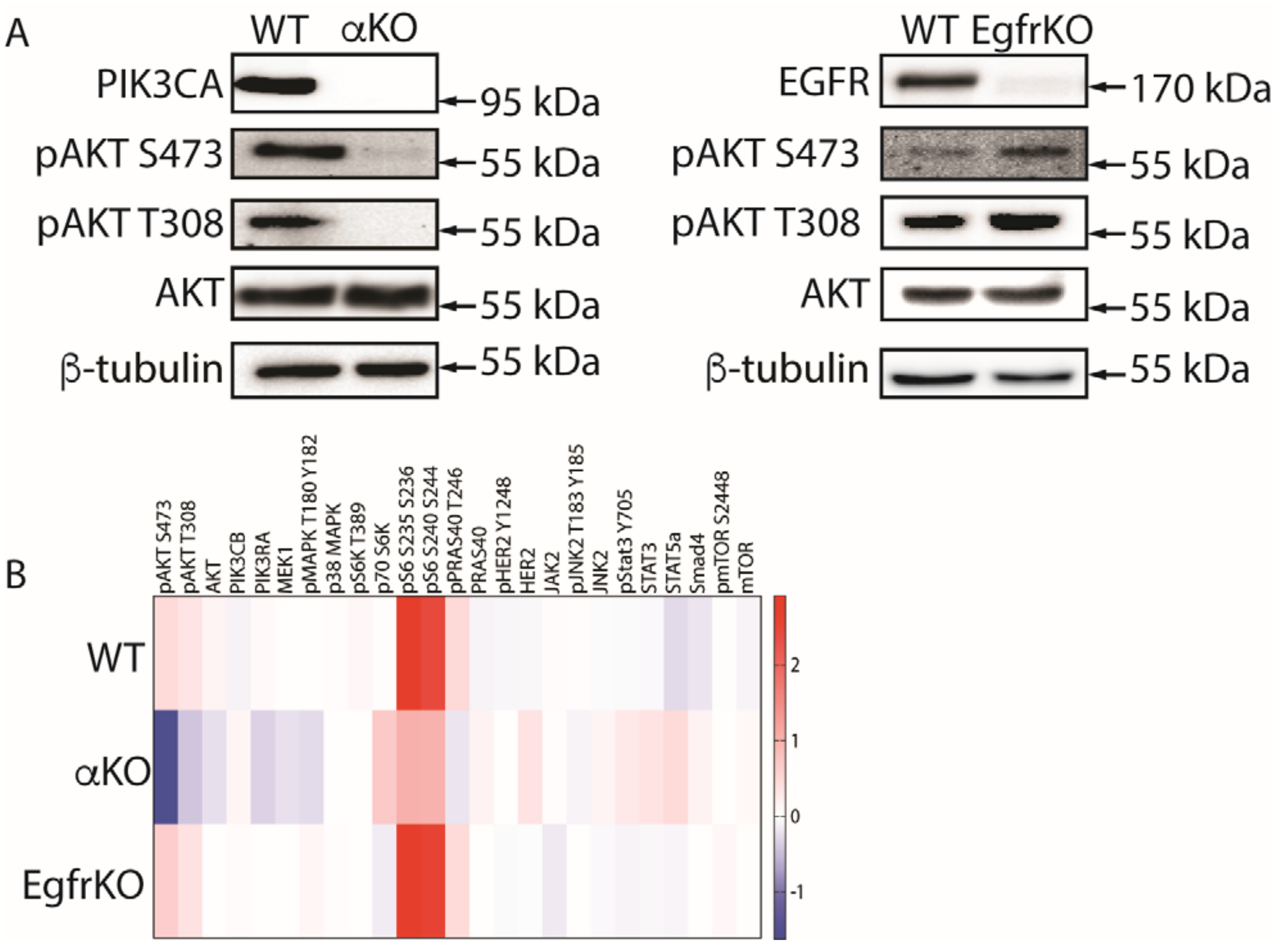


**Supplementary Fig. S2. Western blotting and RPPA analysis of KPC cell lines.** *Pik3ca* or *Egfr* were targeted in wildtype (WT) KPC cells using CRISPR/Cas9 and clonal cell lines were established. **(A)** Representative western blots to confirm gene ablation and to assess AKT activation. β tubulin is a loading control. Experiments were repeated 3 times. **(B)** Heatmap for RPPA analysis of the three cell lines.

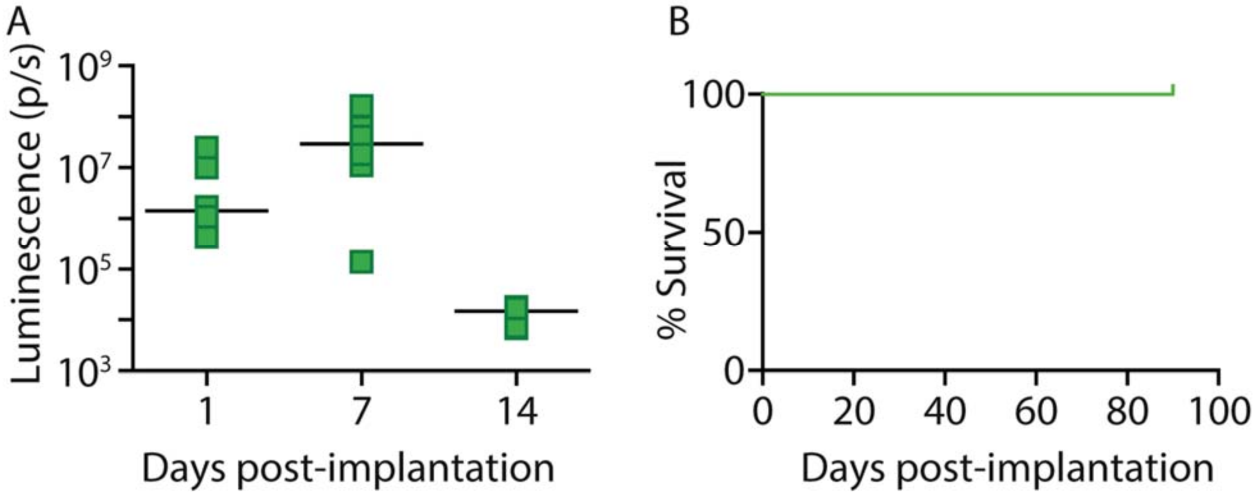


**Supplementary Fig. S3. Regression of αKO clone2 tumors *in vivo*.** αKO clone2 cells (0.5 million) were implanted in the head of the pancreas of B6 mice (*n* = 6) and tumor growth was monitored by IVIS imaging of the luciferase signal. **(A)** Graph shows quantification of luciferase signals. **(B)** Kaplan-Meier survival curve.

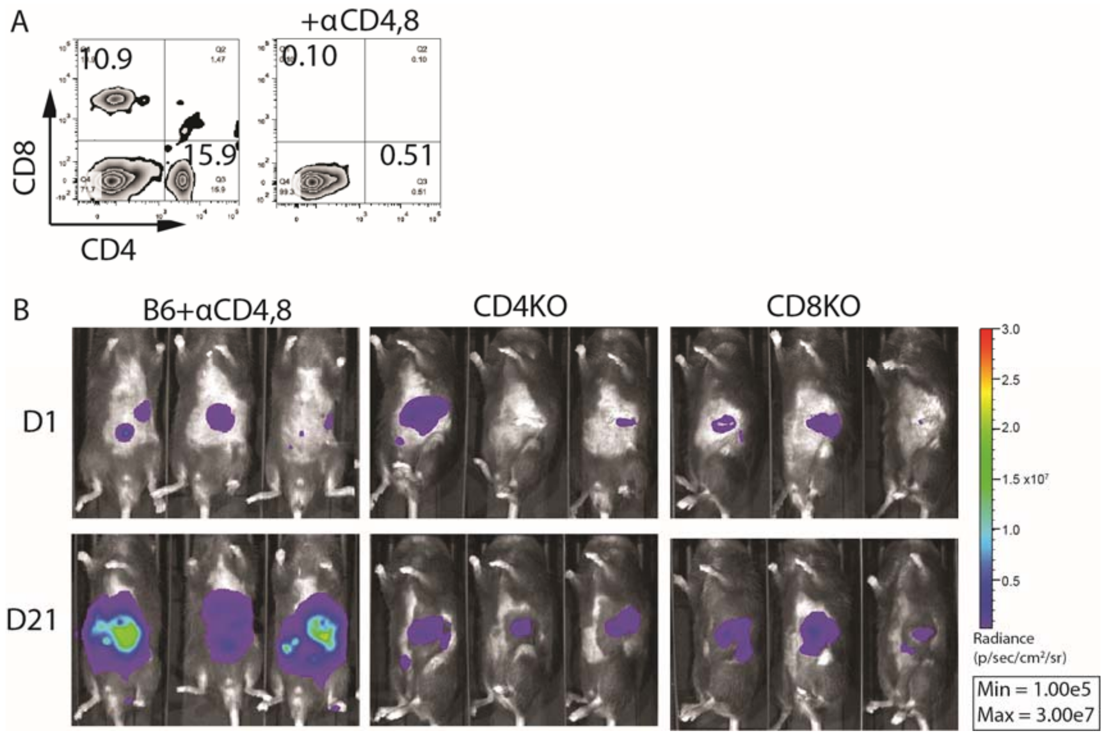


**Supplementary Fig. S4. Tumor growth in mice lacking CD4 and/or CD8 T cells. (A)** B6 mice were injected with neutralizing CD4 and CD8 antibodies. Peripheral blood was analyzed for depletion of $CD4^+$ and $CD8^+$ T cells. Dot plots show CD4 and CD8 T cells in mice injected with saline (*left*) or CD4/8 antibodies (*right*). (**B)** IVIS images of B6 mice injected with CD4/8 antibodies, CD4KO, and CD8KO mice implanted with 0.5 million $\alpha$KO cells in the head of the pancreas.

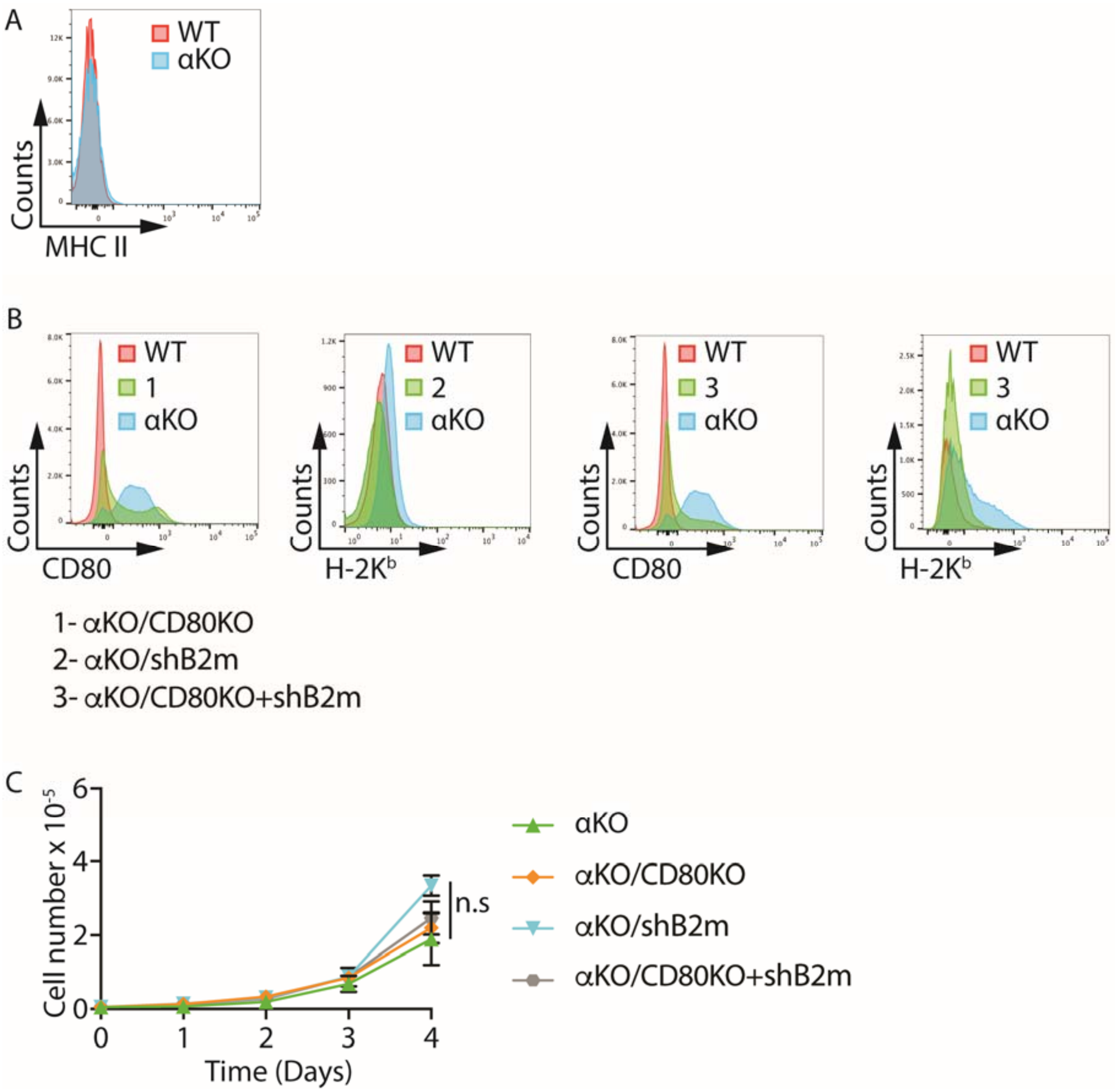


**Supplementary Fig. S5. Cell surface levels of MHC II, H-2K$^b$, and CD80 in KPC cell lines.** Flow cytometric analysis to determine levels of **(A)** MHC II (I-A$^b$) and **(B)** H-2K$^b$ and CD80. **(C)** Proliferation rates of αKO KPC cell lines in standard 2D culture. Cells plated in triplicate were counted at the times indicated (mean ± SEM; *n* = 3). n.s., not significant.

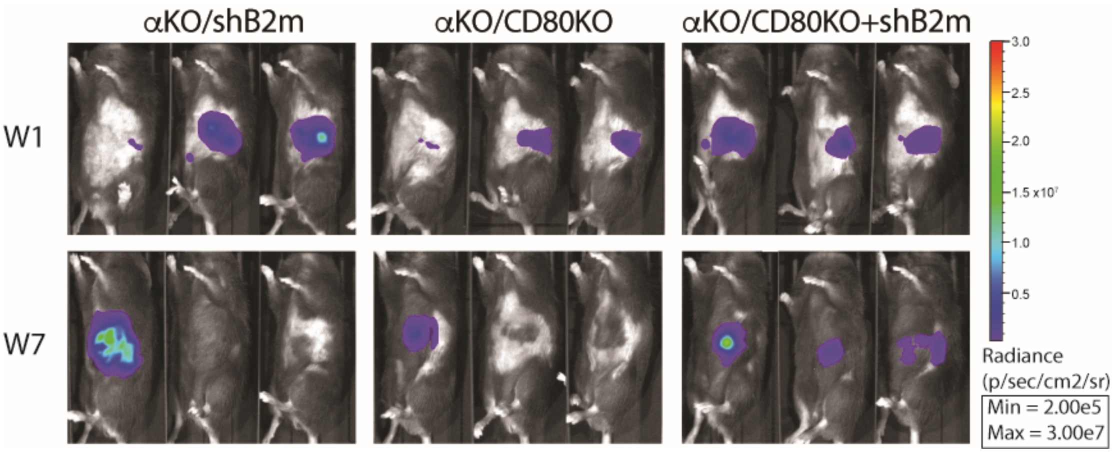


**Supplementary Fig. S6.** ***In vivo* tumor growth.** The indicated cell lines (0.5 million) were implanted in the head of the pancreas of B6 mice. Tumor growth was monitored by IVIS imaging of the luciferase signal. Representative images of 3 mice in each group are shown.

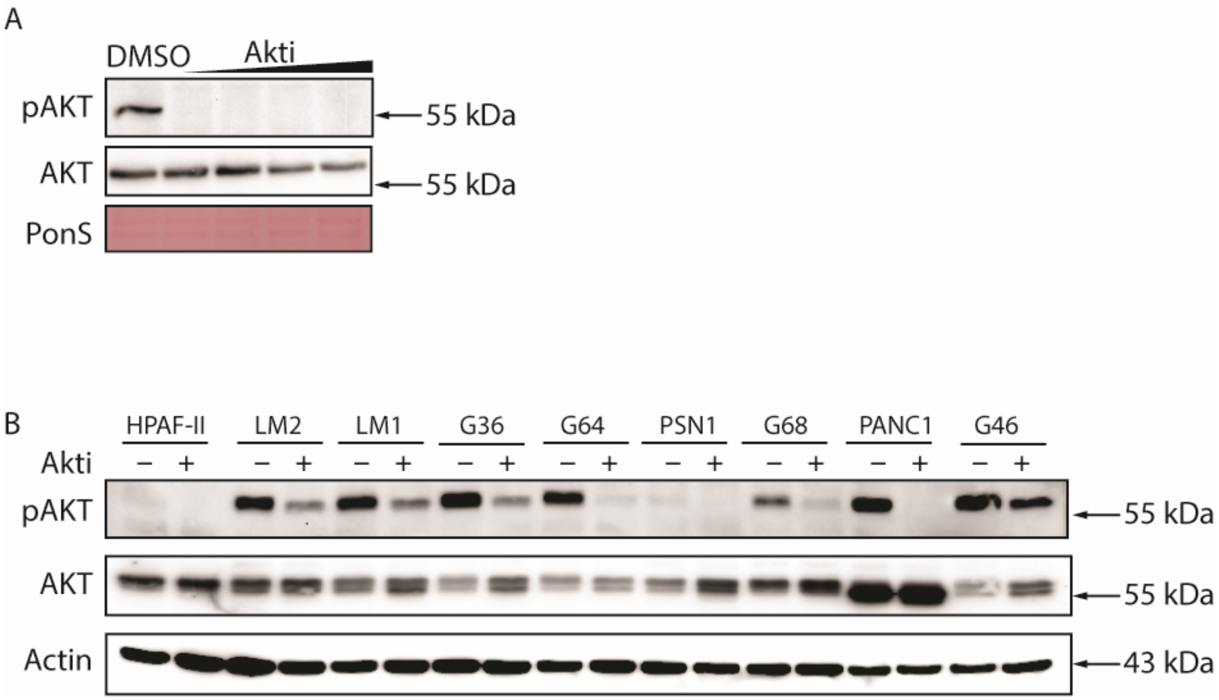


**Supplementary Fig. S7. Effect of Akti treatment on phospho-AKT. (A)** Mouse PDAC cell lines were treated with increasing concentrations of Akti or **(B)** human PDAC cell lines were treated with 10 μM Akti for 48 hours. Cells were lysed in RIPA buffer and total protein was run on a denaturing gel. Western blots show levels of phospho- and total AKT. Actin in a loading control. PonS, Ponceau S-stained blot.

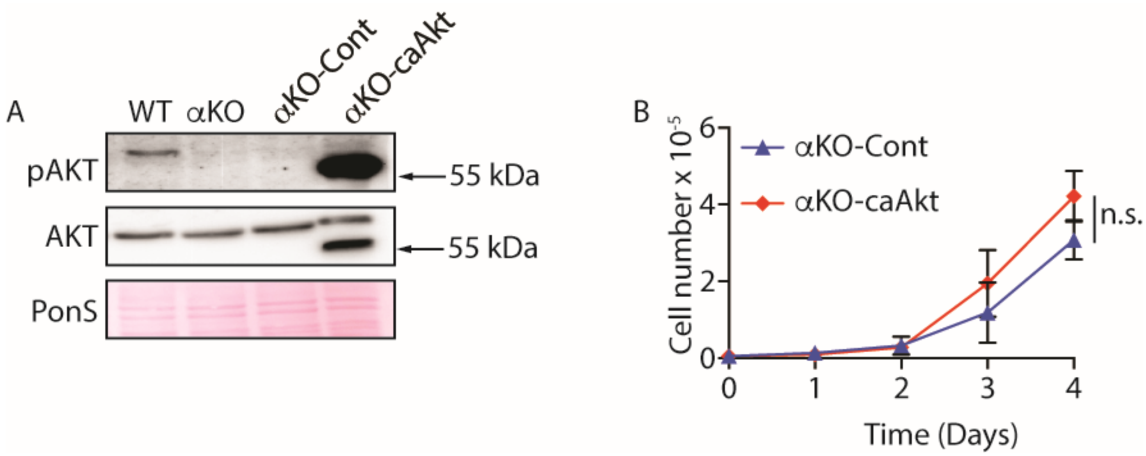


**Supplementary Fig. S8. AKT phosphorylation and *in vitro* growth of αKO-caAkt cells. (A)** αKO cells were infected with lentivirus expressing control vector (Cont) or caAkt. Western blots confirm the expression of caAkt. PonS, Ponceau S-stained blot. **(B)** Proliferation rates in standard 2D culture. Cells plated in triplicate were counted at the times indicated (mean ± SEM; $n = 3$). n.s., not significant.

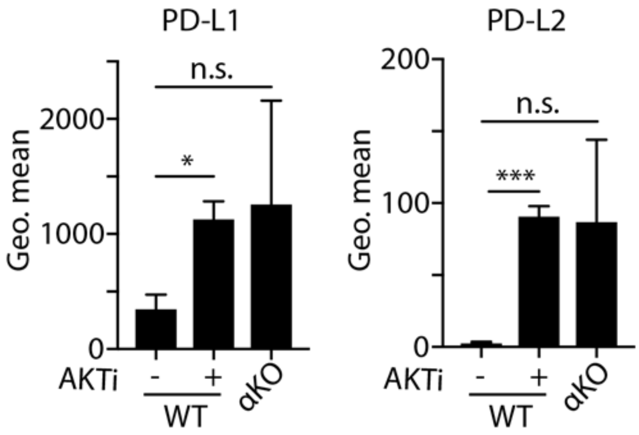


**Supplementary Fig. S9. Cell surface levels of PD-L1 and PD-L2 in WT and αKO cells.** WT cells were treated with 10 μM AKTi for 48 h. Cell surface levels of PD-L1 and PD-L2 were quantified by flow cytometry. WT and αKO cells were used for comparison. The treatment group was normalized to DMSO. The graph shows fold change of geometric means (mean ± SEM) over DMSO. *n* = 4, **P* = 0.0229, ****P* = 0.0008 (paired *t*-test), *n.s.*, not significant.
